# Mapping Risk and Conservation Potential Across the Indo-Pacific with Reefshark Genomescapes

**DOI:** 10.1101/2025.09.22.676358

**Authors:** Shaili Johri, Gonzalo Araujo, John Nevill, Ryan Daly, Theodore EJ Reimer, Zoya Tyabji, Rima W Jabado, Muhammad Ichsan, Iqbal Sani, Robert Schallert, Daivd Curnick, Chi-Ju Yu, Barbara A Block

## Abstract

Overfishing has severely depleted marine populations worldwide, including within protected areas. Illegal and unreported fishing are major contributors to this decline. Large-bodied apex predators such as sharks are among the most affected, with overfishing causing dramatic species declines and ecosystem destabilization due to trophic downgrading. Key barriers to effective marine conservation and management include: Data deficiencies that hinder population benchmarks and impact assessments, limited surveillance, allowing illegal fisheries to disproportionately affect apex predators, and insufficient capacity in vulnerable nations to monitor and protect species within their waters.

Our study addresses these challenges through a novel genomic framework that enables assessment of shark population diversity and health, while also improving fisheries traceability by detecting instances of illegal fishing across the Indian and Pacific Oceans. We present the *Reefshark Genomescape*, the first genome-wide reference database for Indo-Pacific reef sharks, an assessment of genetic diversity, structure, and connectivity of two key species across their Indo-Pacific range and geographic assignment of fished individuals using population-specific genetic signatures.

We show that grey reef shark (*Carcharhinus amblyrhynchos*) populations exhibit high genetic diversity, strong population structure, and elevated F_st_ values, with previously unknown connectivity between the central and western Indian Ocean and clear isolation of populations in the Andaman Sea. In contrast, silvertip sharks (*Carcharhinus albimarginatus*) display high connectivity, but show genomic signals of declining population health, supporting a reassessment of their IUCN status. Using supervised machine learning with Monte Carlo cross-validation, we assigned geographic origins to fished grey reef sharks with 96% accuracy.

These findings provide critical insights into population structure, connectivity, and health of two ecologically important reef shark species, while establishing a robust method for assigning geographic origin. We anticipate this framework will support regional conservation assessments and targeted management. Moreover, by enabling the identification of fishing hotspots and detection of IUU fishing, it lays the groundwork for a broader traceability system in marine ecosystems. Much like the landmark elephant ivory tracing study, our approach has the potential to transform marine conservation globally.

**Graphical Abstract:** We developed the Reefshark Genomescape, a genomic framework for assessing shark population health and fisheries traceability across the Indo-Pacific. Genome-wide data from grey reef and silvertip sharks revealed contrasting patterns, unexpected connectivity, and genomic signals of decline. Geographic assignment of fished individuals reached 96% accuracy, enabling detection of illegal fishing and identification of hotspots. This framework strengthens regional management, supports IUCN reassessments, and lays the foundation for global marine traceability systems.

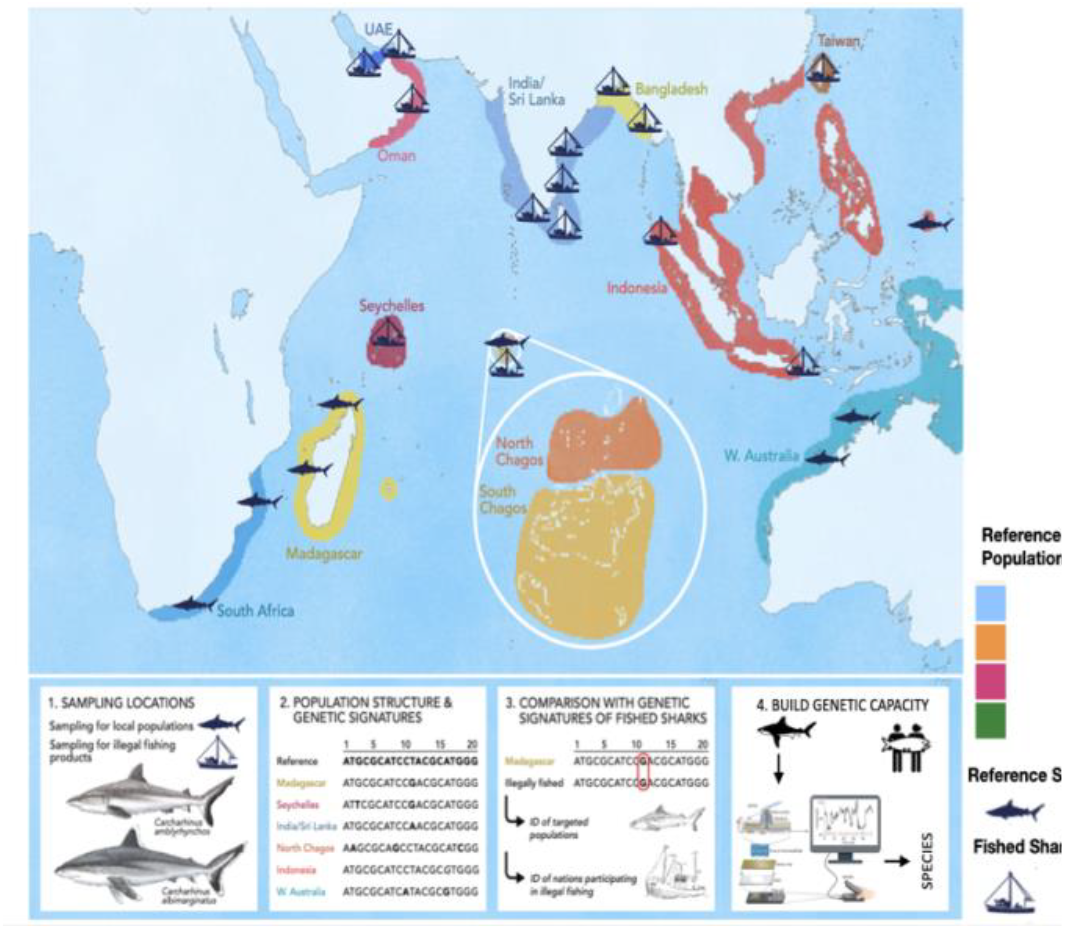

## INTRODUCTION

Marine ecosystems across the globe are experiencing unprecedented biodiversity loss due to overfishing and exploitation, driven by the rising global demand for seafood. Coral reefs, one of the most biodiverse ecosystems on earth, and home to over one-third of all marine fish species (Plaisance et al. 2011) are especially vulnerable to overfishing. This has contributed significantly to reef degradation by direct removal of key predator species (Sherman et al. 2023; Dulvy et al. 2021) and the indirect disruption of ecological dynamics once they are gone (Graham et al. 2017; Sandin et al. 2008). Illegal, unreported, and unregulated (IUU) fishing is a major driver contributing to overfishing in reef ecosystems especially in East and West Africa, North Pacific and Asia, which are frequent hotspots for IUU fisheries (Belhabib and Le Billon 2020).

The threats faced by reefs often vary spatially, necessitating region-specific conservation strategies (Pazmiño et al. 2018). However, due to technical, legal, and geopolitical loopholes in vessel monitoring systems (Paolo et al. 2024) and as a result of extensive IUU fishing activity, it is challenging to assess where fishing efforts are focused, which species and stock populations are being depleted and which reef ecosystems need bolstered protections. Many studies have identified species targeted by reef fisheries (Fields et al. 2015), and some have examined specific stock populations (Fields et al. 2020). A very few studies assess ecosystem health, genetic diversity, evolutionary capacity, effective population size, and sustainable yields at a trans-oceanic scale. This knowledge gap constrains long-term conservation planning that prioritizes species and ecosystems by urgency and adapts to shifting distributions and fishing thresholds under climate and anthropogenic pressures.

Reef sharks comprise a significant portion of the large predator biomass in coral reef ecosystems (Sandin et al. 2008; Friedlander et al. 2014) and are among the primary targets of coral reef fisheries (Graham et al. 2017). As a result, reef shark populations have declined sharply worldwide due to overfishing and exploitation (Sherman et al. 2023). Many reef-associated shark species, display strong site fidelity and limited gene flow across distant reef systems, resulting in genetically distinct populations (Pazmiño et al. 2017; Momigliano et al. 2017; Maisano Delser et al. 2019). While this population structure increases vulnerability to local extinctions, it enables the development of genomic reference libraries that identify the geographic origin of individuals and establish population-specific baselines. As apex and mesopredators, reef sharks serve as key indicators of coral reef health and fishing pressure. Population-specific genomic studies thus provide powerful tools for monitoring both reef shark populations and overall ecosystem health, informing conservation at ecosystem scales. Here we use whole genome resequencing to identify population structure across the Indo-Pacific basin in grey reef sharks (*Carcharhinus amblyrhynchos*) and silver tip sharks (*Carcharhinus albimarginatus*) and to develop tools for identification of geographic origin of shark catches within fisheries.

Grey reef sharks are dominant and ecologically important predators across much of the tropical Indo-Pacific reef ecosystems, serving as key indicators of ecosystem integrity (Friedlander et al. 2014). Despite their ecological importance, they are listed as Endangered on the IUCN Red List, with an estimated global biomass decline of 50–79% (Ali et al. 2020). Studies of movement patterns and long-term reef fidelity show that grey reef sharks have small activity spaces, averaging 58 km in radius, have low transboundary crossings and exhibit high reef residency (Espinoza et al. 2014; Jacoby et al. 2020; Daly et al. 2023; Dwyer et al. 2020). These movement and fidelity patterns, along with regional genetic studies in Australia and Indonesia (Momigliano et al. 2017), indicate strong population structuring. Combined with the species’ sharp population declines, these findings underscore the need for population-specific assessments to inform global conservation status and guide fishing and trade regulations.

In contrast, silvertip sharks have a patchy distribution across the Indo-Pacific but are among the most frequently observed reef sharks within their range (Bond et al. 2015). They are classified as Vulnerable, with a predicted global biomass decline of ~30% over the past 54 years (Rigby, C.L. et al. 2023). Silvertip sharks are more wide-ranging and dynamic than grey reef sharks, with activity spaces averaging 175 km— nearly three times larger, and lower site fidelity (Jacoby et al. 2020). Regional genetic studies using reduced-representation markers indicate high connectivity within the southwest Pacific but isolation between populations separated by open ocean (Green et al. 2019). These patterns highlight the need to delineate management units and identify key ocean corridors used by silvertip sharks to ensure effective population protection.

Both species are heavily impacted by fisheries and illegal trade across their Indo-Pacific range (Dulvy et al. 2014; Dulvy et al. 2021; Bennett et al. 2022), and are listed under Appendix II of the Convention on International Trade in Endangered Species of Wild Flora and Fauna (CITES); yet enforcement of international trade regulations remains a challenge (Bond et al. 2025). Their broad but discontinuous distributions and ongoing population declines create an urgent need for fine-scale, spatially explicit assessments of genetic connectivity, population structure, and demographic health across their ranges. While regional studies have provided important insights into local connectivity patterns, no study to date has examined genome-wide population structure of either species across the full Indo-Pacific.

In addition to assessing conservation status, the contrasting movement and connectivity patterns of grey reef and silvertip sharks offer a unique opportunity to study population connectivity and structure across Indo-Pacific reefs. Developing a genetic reference database spanning their distributions will answer key ecological questions about how ecological, oceanographic, and historical factors shape connectivity, and will establish population-specific baselines. Matching fished individuals to this database can provide valuable insights into spatial fishing pressures, identify fishing hotspots, and detect instances of IUU fishing and trade. Both species show strong potential as biological indicators, and this approach can transform fisheries monitoring, improve stock assessments, and support more effective, adaptive conservation and management of coral reef ecosystems. Importantly, these efforts not only enable targeted protection of reef shark populations but also contribute to the broader conservation of reef ecosystems by detecting spatially explicit fishing pressures.

In this study, we present a genomic framework that both assesses the demographic health and connectivity of reef shark populations and supports fisheries traceability and conservation enforcement. We constructed a comprehensive genetic reference database for two key reef shark species, *Carcharhinus amblyrhynchos* and *C. albimarginatus*, which also serve as indicators of coral reef ecosystem health. By collecting samples across their Indo-Pacific distributions, we conducted the first range-wide population genomic analysis of these species, integrating complete mitochondrial genomes with hundreds of thousands to millions of genome-wide single nucleotide polymorphisms (SNPs).

Our dataset spans multiple regions, including the western Indian Ocean, Chagos Archipelago, Andaman Sea, eastern Indian Ocean, and the broader Pacific. These regions differ in reef density, geographic isolation, and exposure to fishing pressures, providing an ideal framework to investigate how ecological and anthropogenic factors shape genetic structure and gene flow. Using this approach, we address critical gaps in understanding reef shark population connectivity and resilience to depletions and demonstrate how genomic data can inform fisheries traceability, CITES reporting, and sustainable marine resource management. In doing so, this study provides a baseline health and diversity index for two ecologically important reef predators across their Indo-Pacific population range. The data and analytical tools presented provide a framework for evaluating regional conservation status, fishing pressures, and guiding targeted effective management strategies.

## METHODS

### Biological Materials

To obtain reference samples fin clips were taken during catch and release operations while tagging grey reef and silvertip sharks across sampling locations. Sharks were captured using single monitored drum lines with barbless circle hooks (14/0−18/0) baited with tuna (Scombridae spp.). On capture, sharks were inverted and restrained submerged alongside a research vessel. Individuals were sexed (presence/absence of claspers) and measured (pre-caudal length and total length, in cm) before taking a genetic sample from the trailing edge of the anal fin. Sharks were restrained for less than ten minutes and were released at the site of capture.

To obtain reference samples from fished sharks, fin clips or muscle tissue were collected from artisanal fishers or distant fishing fleets at landing sites, fish markets, or directly on fishing vessels. Samples were stored immediately after collection in RNA later tubes or in 90% ethanol at room temperature and then stored once ethanol was drained off in a −80 C freezer until DNA isolation. Samples were collected at the following locations indicated in Figure 1. The number of samples per site is listed in Table 1.

**Table 1:**
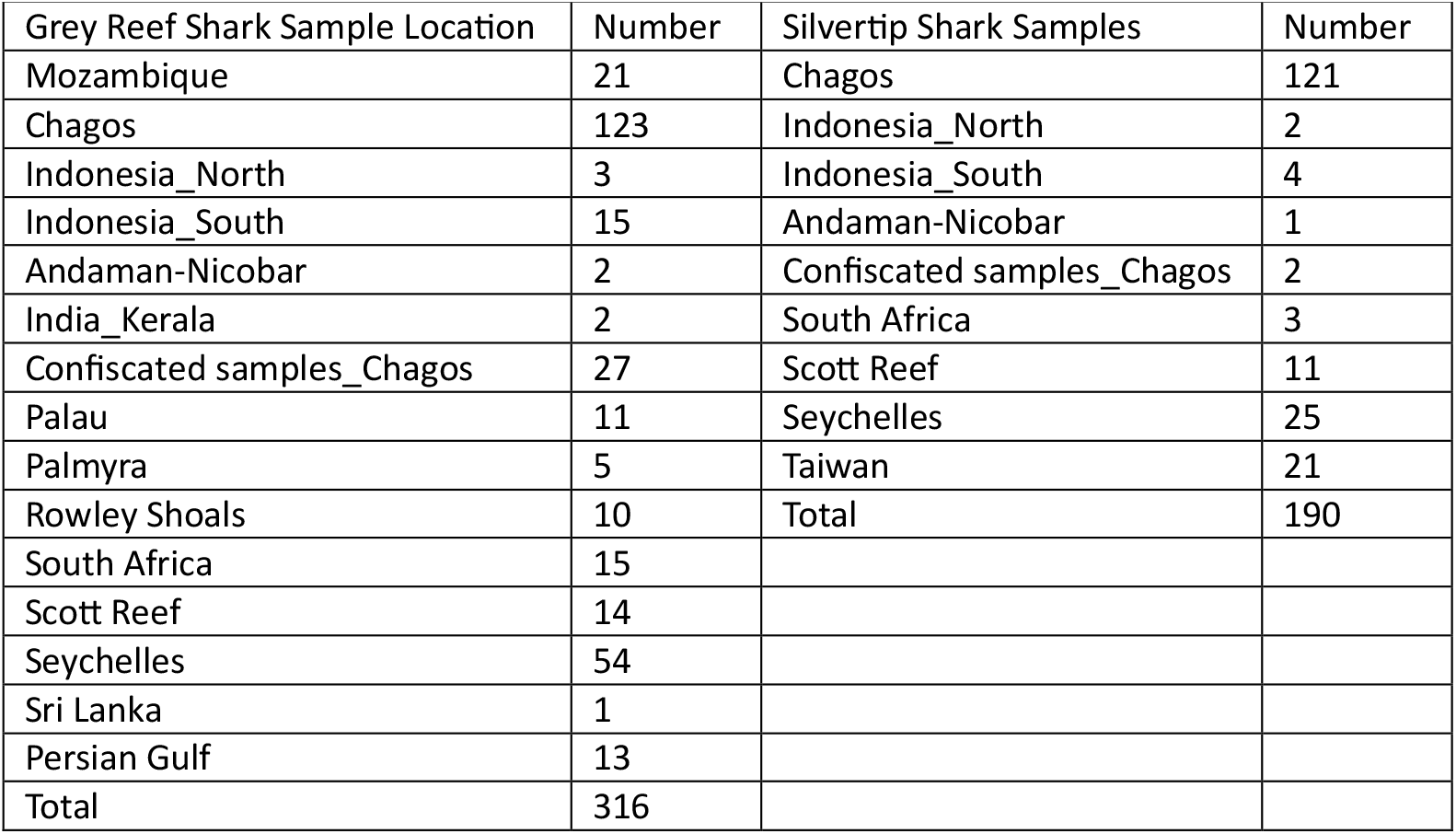
Sampling Locations and number of samples.

| Grey Reef Shark Sample Location | Number | Silvertip Shark Samples | Number |
| --- | --- | --- | --- |
| Mozambique | 21 | Chagos | 121 |
| Chagos | 123 | Indonesia_North | 2 |
| Indonesia_North | 3 | Indonesia_South | 4 |
| Indonesia_South | 15 | Andaman-Nicobar | 1 |
| Andaman-Nicobar | 2 | Confiscated samples_Chagos | 2 |
| India_Kerala | 2 | South Africa | 3 |
| Confiscated samples_Chagos | 27 | Scott Reef | 11 |
| Palau | 11 | Seychelles | 25 |
| Palmyra | 5 | Taiwan | 21 |
| Rowley Shoals | 10 | Total | 190 |
| South Africa | 15 |  |  |
| Scott Reef | 14 |  |  |
| Seychelles | 54 |  |  |
| Sri Lanka | 1 |  |  |
| Persian Gulf | 13 |  |  |
| Total | 316 |  |  |

**Figure 1.**
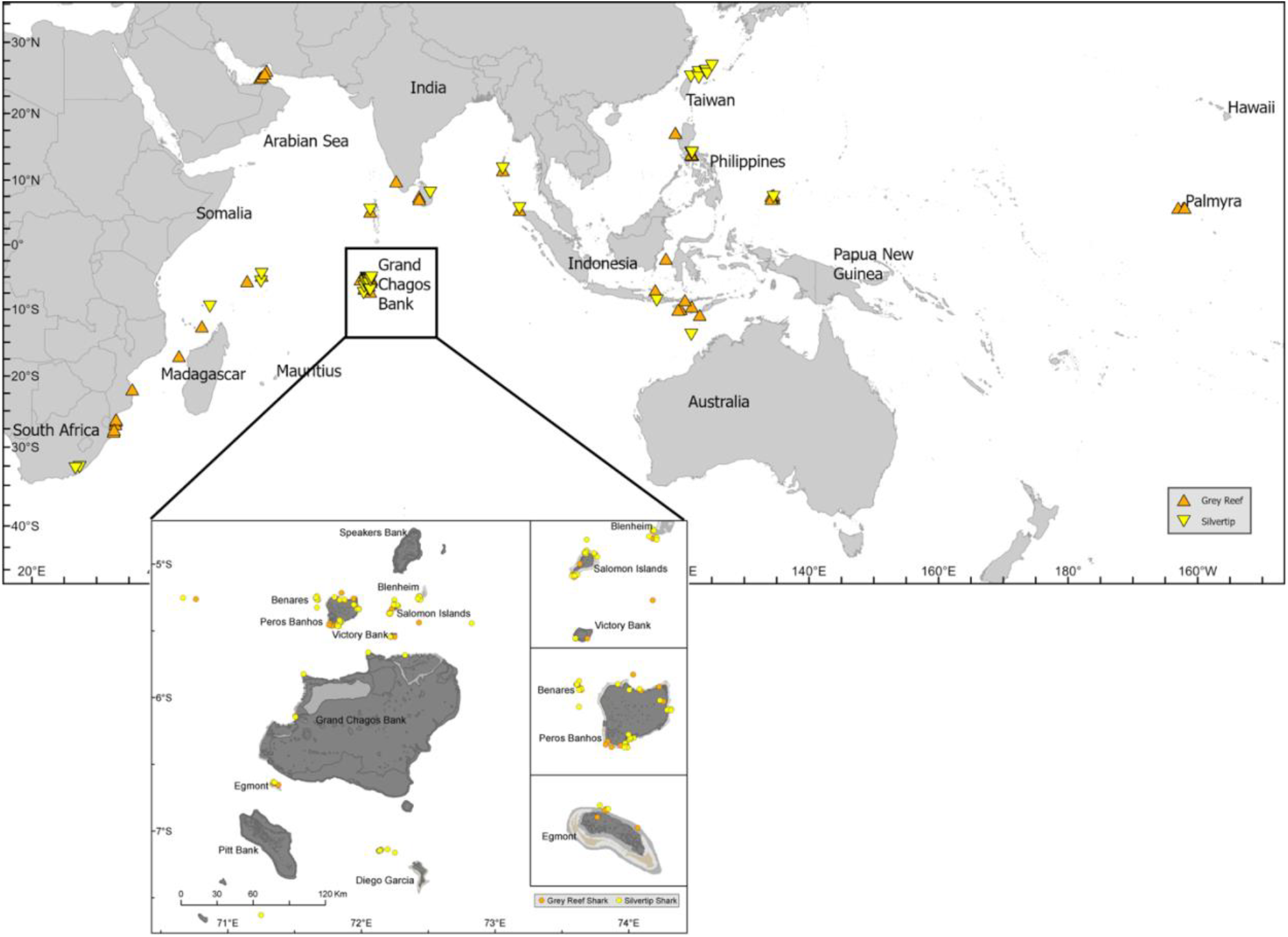
Map showing collection sites of the grey reef (orange triangles) and silver tip (yellow triangles) shark samples.

### DNA extraction and library preparation

Each fin clip sample was subsampled and DNA was extracted from a 10 mg piece using the DNeasy Blood & Tissue Kit (Qiagen N.V., Venlo, The Netherlands), including a 3 hour lysis step. The concentration and quality of genomic DNA was checked using a NanoDrop ND-1000 Spectrophotometer (Thermo Scientific, Inc., Waltham, MA, USA) and an Agilent Tape Station 4150 (Agilent Technologies, Inc., Santa Clara, USA). For most of the samples DNA was diluted to be within 10-100ng/ul and sent for Library Preparation and sequencing to Genomics & Bioinformatics Services, College Station, TX, USA.

Libraries were prepared using the PerkinElmer NextFlex DNA Library Prep Kit for Illumina (Perkin Elmer Inc., Connecticut, USA). Library quality and quantity was checked on an Agilent Tape Station 4150 (Agilent Technologies, Inc., Santa Clara, USA). Sequencing was performed on the Illumina NovaSeq 6000 S4 × 2×150 v1.5 platform (Illumina Inc., San Diego, USA) at the sequencing core Genomics & Bioinformatics Services, College Station, TX, USA. Samples from India were processed and sequenced using the same methods and platform at Clevergene Inc., Bangalore, India.

### Mitochondrial DNA Analyses

Whole mitochondrial genomes were generated from the trimmed paired whole genome sequence reads of 316 grey reef sharks and 190 silvertip sharks using the software package Mitos and MitoFinder (Bernt et al. 2013). Annotated coding and non-coding regions for each mitogenome were assembled into mitochondrial contigs and aligned to the reference mitogenome (Dunn et al. 2020; Johri et al. 2020) for the respective species to generate a multi sample nucleotide alignment file using Geneious^R^ software. Diversity indices (number of haplotypes NH), within population diversity and measures of pairwise genetic differentiation (Φ_ST_) were estimated in the software Popart v. 1.7 (Leigh and Bryant 2015). A minimum spanning network of whole mitochondrial genome haplotypes was constructed in the software Popart v. 1.7 (Leigh and Bryant 2015).

### Whole Genome Data Processing

Adapters, low quality bases and 5’ and 3’ start or end sequences were trimmed with Trimmomatic v.0.39 (Bolger et al. 2014). Cleaned reads were then mapped to the respective grey reef or silvertip shark reference genomes (NCBI accession number JBNGOJ000000000 and JBNGOK000000000, (Johri et al. 2025, manuscript in review) using BWA-MEM v.0.7.17-r1188 (Li and Durbin 2009). Reads were sorted by query name and duplicates were removed via Picard v. 2.26.6 (“Picard Tools - By Broad Institute,” n.d.). Sorted bam files for 316 samples for the grey reef shark and 190 samples for the silvertip shark were combined into two vcf files respectively using BCFtools mpile-up (Li et al. 2009). VCFtools was used for downstream SNP quality control which included filtering the VCF files for: a minimum base call and mapping quality threshold of 30, only reads mapping to a single location of the genome retained, Minor Allele Frequencies (MAF) > 1%, Hardy–Weinberg Equilibrium (HWE), read depth of 3X, biallelic loci only, and a call rate threshold > 75% (i.e. a total of 25% missing data) (Danecek et al. 2011). Finally the VCF file was filtered for linked loci using Plink at linkage disequilibrium (LD) r2 > 0.6 (Chang 2020).

### Analysis of genetic structure and diversity

A relatedness analysis was performed using VCFtools to prevent biased intra-population diversity results. In order to test for genetic homogeneity between locations we calculated pairwise fixation indices, F_ST_ for nuclear SNP markers using the R package ‘StAMPP’ (https://cran.r-project.org/web/packages/StAMPP) (Pembleton et al. 2013). Each analysis consisted of 100,000 bootstraps generating confidence intervals and P-values for each pairwise comparison of F_ST_. Significance levels of all pairwise tests were corrected for multiple comparisons with a sequential Bonferroni procedure (BF P ¼ conventional P-value 0.05 divided by the number of tests per marker type; Rice 1989).

The VCF file was converted to BED format and Principal Component Analyses conducted using Plink2. Using the same BED files, admixture proportions were estimated using the Admixture software package (Alexander et al. 2009) for k = 2-6 for both the grey reef shark and silvertip shark populations. The cross validation (CV) error was estimated for each of the K=2-6 and the Ks with lowest CV error was used for subsequent admixture analyses of both species’ population groups. The PCA and Admixture plots were generated using the eigenvec and eigenval files and the.Q and.P files respectively in RStudio.

Gene flow patterns were examined under the assumption of asymmetric bidirectional gene flow, using the Genepop file with a set of 11,345 SNPs across all locations and 316 individuals for the grey reef shark and 190 individuals for the silvertip shark. This analysis was executed using the R-function divMigrate from the diveRsity R-package (Keenan et al. 2013). The divMigrate function computes migration rates between all sites and subsequently normalizes them, generating relative migration rates from zero to one. Furthermore, significant asymmetry in gene flow direction between two sites was assessed employing statistics introduced by Alcala et al. This approach combines information from both genetic differentiation measures, GST and D, to estimate the number of migrants per generation Nm, from which relative migration is derived. To evaluate the presence of significant asymmetric migration, 1,000 bootstraps were conducted.

Nucleotide diversity was estimated by generating a VCF file using BCFtools that contained variant and non-variant sites for each of the grey reef and silvertip shark populations. The VCF file thus generated along with the population data for each sample was used to calculate nucleotide diversity per population using the software package pixy v0.95.02 (Korunes and Samuk 2021).

### Demographic History

Demographic histories of the grey reef shark and the silvertip shark were reconstructed using Pairwise Sequential Markovian Coalescence (PSMC) v0.6.5 (Schiffels and Durbin 2014). The Illumina sequence reads from each species were mapped to the respective chromosomal scaffolds for the species using BWA-MEM v0.7.17 (Li and Durbin 2009) and the consensus sequence was called using BCFtools v1.21-79-gcef68bc+ mpileup (Li et al. 2009), BCFtools call, and vcfutils “vcf2fastq.” function in the PSMC software package (https://github.com/lh3/psmc/utils). Minimum depth cutoffs of 5 and maximum depth cutoffs of 100 were applied to all genomes using vcfutils. Remaining positions with an average mapping quality greater than 30 were used to create the input file for PSMC. The program `fq2psmcfa’ in the PSMC software package, was used to transform the consensus sequence into a fasta-like format. In order to perform bootstrapping, we ran ‘splitfa’ function in PSMC, to split long chromosome sequences to shorter segments. PSMC was run for 25 iterations with parameter values: -N25 -t15 -r5 -p “4 + 25*2 + 4+6”. We then performed 100 rounds of bootstrapping to evaluate variance in effective population size (*N*_*E*_) estimates. Results from the 100 rounds were concatenated into a combined PSMC file, which was then used to generate plots using ‘psmc_plot.pl’ from the PSMC package. In order to visualize the PSMC plot, we applied a mutation rate of 1.9e−08 (Lesturgie et al. 2022) and a generation time of 7-10 years for the grey reef shark (Bm et al. 1997; Lesturgie et al. 2022). For the silvertip shark we applied the mutation rate 1.9e−08, same as the grey reef shark as has been used previously for multiple reef shark species (Lesturgie et al. 2022). A generation time of 9.5 years was used for the silver tip shark (Kato, S. and Hernandez 1967).

### Heterozygosity and Runs of Homozygosity

After self-alignment of Illumina short reads from the same individual to the reference genome, heterozygosity estimates were calculated using ANGSD v.940 (Korneliussen et al. 2014). Heterozygosity estimates for the whole genome and on a per site basis were obtained using these methods. We calculated homozygosity for the two genomes by aligning short reads from the same individual to the genome and assessing variant and non-variant sites across the genome. Short reads were mapped to the genome using BWA-MEM v0.7.17 (Li and Durbin 2009) and BCFtools mpile-up (Li et al. 2009; Danecek et al. 2021) was used to generate a VCF file. BCFtools ROH (Narasimhan et al. 2016) was used to identify runs of homozygosity across the genome. Illumina short read data at 30x or higher genome coverage was used to create a vcf file which was then used to determine ROH length and location across the two genomes using BCFtools ROH.

### Population Assignment of Fished Sharks

For the population assignment of fished shark specimens with unknown source populations we generated a supervised machine learning model using Monte Carlo Cross Validation in the Rpackage assignPOP (Chen et al. 2018). Monte-Carlo cross-validation was conducted using 75%, 80% and 95% of individuals from each population by the top 10%, 25%, 50% of high F_ST_ and all loci as training sets and each training set combination was resampled for 30 iterations. A total of 360 assignment tests were performed. Support vector machine (model = “svm”) classification model was used to build predictive models. Assignment accuracy of the Monte–Carlo cross-validation results was calculated and the accuracy plot generated. After an optimal model with high assignment accuracy of >95% for the training dataset was generated, the model was used to assign populations for fished shark specimens. Each of the svm, naïveBayes, random forest or lda models were tested to the best assignment accuracy of unknown samples.

## RESULTS

### Genome-wide SNP Analysis in *Carcharhinus amblyrhynchos*

Principal component analysis (PCA) of 6 million genome-wide SNPs from 323 *C. amblyrhynchos* individuals identified four distinct genetic clusters and one intermediate group. The intermediate cluster consisted of two individuals from northern Indonesia, likely reflecting recent admixture between the Andaman and Chagos populations (Figure 2A). These four clusters were further supported by admixture analysis (Figure 2B). Cross-validation errors were lowest for K = 2, 3, and 4, with K = 4 providing the most informative resolution of population structure of *C. amblyrhynchos*.

**Figure 2.**
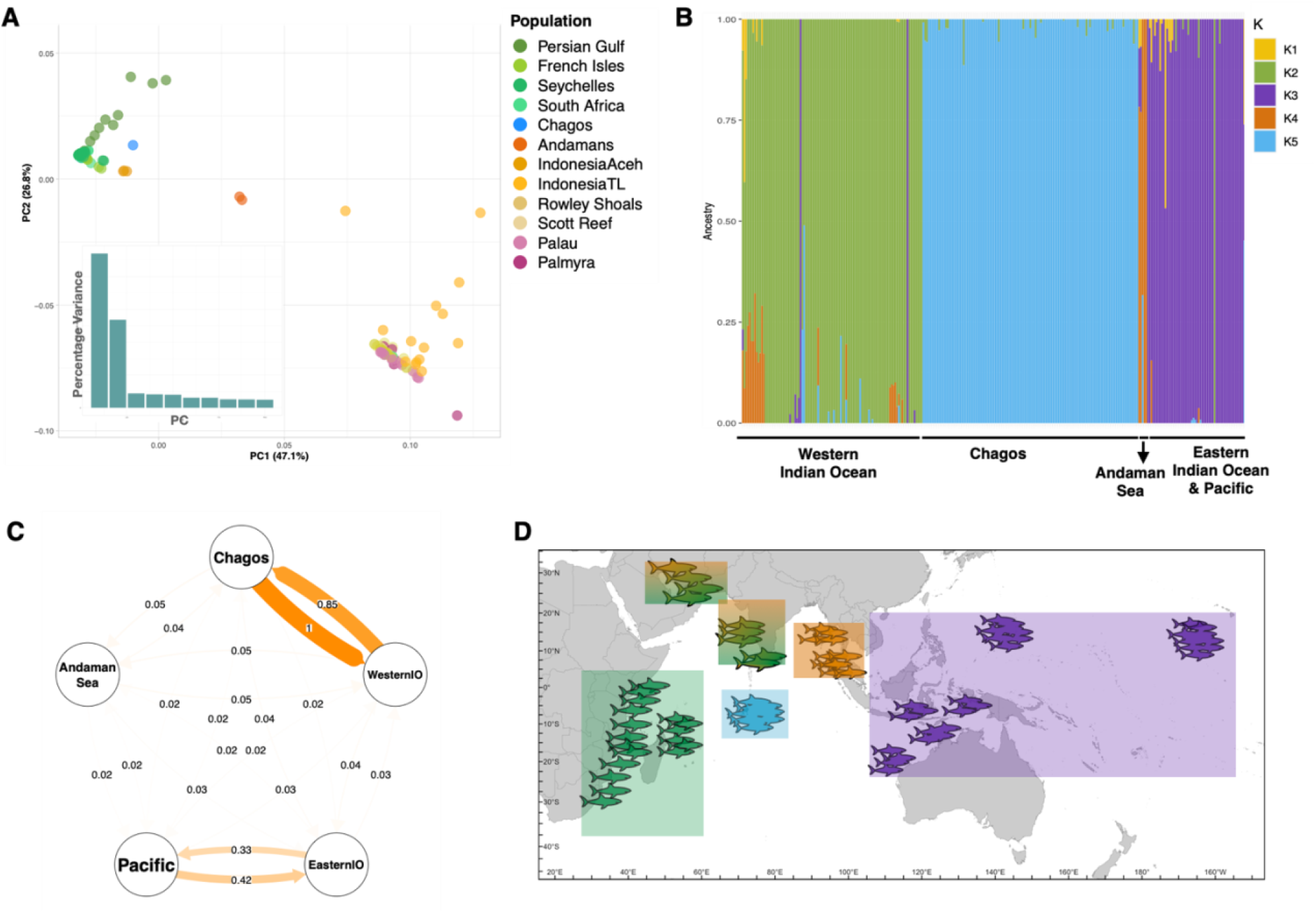
Population structure in grey reef sharks across the Indo-Pacific. (A) Principal component analysis (PCA) showing clear genetic differentiation among regional groups. (B) Admixture plot highlighting ancestry proportions and levels of genetic mixing between populations. (C) Directional migration and gene flow plot, illustrating asymmetrical connectivity among regions. (D) Map of resulting sub-populations defined from these analyses, showing four major genetic clusters (WIO, Chagos, Andaman, and EIO–Pacific) that represent distinct management units.

At K = 4, four well-differentiated clusters were resolved: (i) the WIO, (ii) Chagos archipelago, (iii) the Andaman–northern Indonesia group, and (iv) the EIO–Pacific group, which included southern Indonesia, Western Australia, Palau, and Palmyra (Figure 2B). At K = 2, two major genetic clusters were recovered: one comprising the EIO and Pacific populations, and the other including all WIO populations along with the Chagos archipelago (Supplementary Figure 1). At K = 3, the Chagos archipelago population emerged as a distinct genetic cluster (Supplementary Figure 1).

Genome-wide F_ST_ analyses between individual populations and regional clusters, confirmed strong genetic differentiation consistent with PCA and admixture results (Supplementary Figure 2A, Supplementary Table 1 & 2). The Chagos population showed the highest differentiation from the Pacific (F_ST_ = 0.284), followed by the EIO cluster (F_ST_ = 0.242), moderate differentiation from the Andaman Sea (F_ST_ = 0.107), and the lowest from the WIO (F_ST_ = 0.007). The smallest differences were between the Eastern IO and Pacific (F_ST_ = 0.008) and between the WIO and Chagos (F_ST_ = 0.007).

Pairwise FST values between individual populations mirrored regional trends (Supplementary Figure 2B). Populations within the same regional cluster showed low genetic differentiation, while FST values increased with geographic distances. The WIO populations were most differentiated from those in the Andaman and EIO clusters. Notably, the Chagos population exhibited greater genetic distance from Andaman and EIO populations than from the WIO, despite geographic proximity to the Andamans. Similarly, populations within the Andaman cluster were highly differentiated from those in the EIO and broader Indo-Pacific, underscoring pronounced genetic structuring over relatively short distances.

To assess gene flow among regions, we analyzed 10,000 high F_ST_ SNPs from 293 individuals, using 1,000 bootstrap replicates and assuming asymmetric bidirectional migration. Results showed significantly high gene flow between Chagos and the WIO, low gene flow between the EIO and Pacific, and no detectable gene flow between any other regions (Figure 2C). The resulting population group assignments, reflecting the major genetic clusters identified for the grey reef sharks, are shown in Figure 2D.

### Genome-wide SNP Analysis in *Carcharhinus albimarginatus*

PCA of 500,000 SNPs from 195 *C. albimarginatus* individuals revealed three main genetic clusters corresponding to the WIO, Chagos, and the EIO–Indo-Pacific region, with some individuals exhibiting intermediate genotypes (Figure 3A). Admixture analysis refined these patterns (Figure 3B), with both K = 2 and K = 3 showing similarly low cross-validation errors.

**Figure 3.**
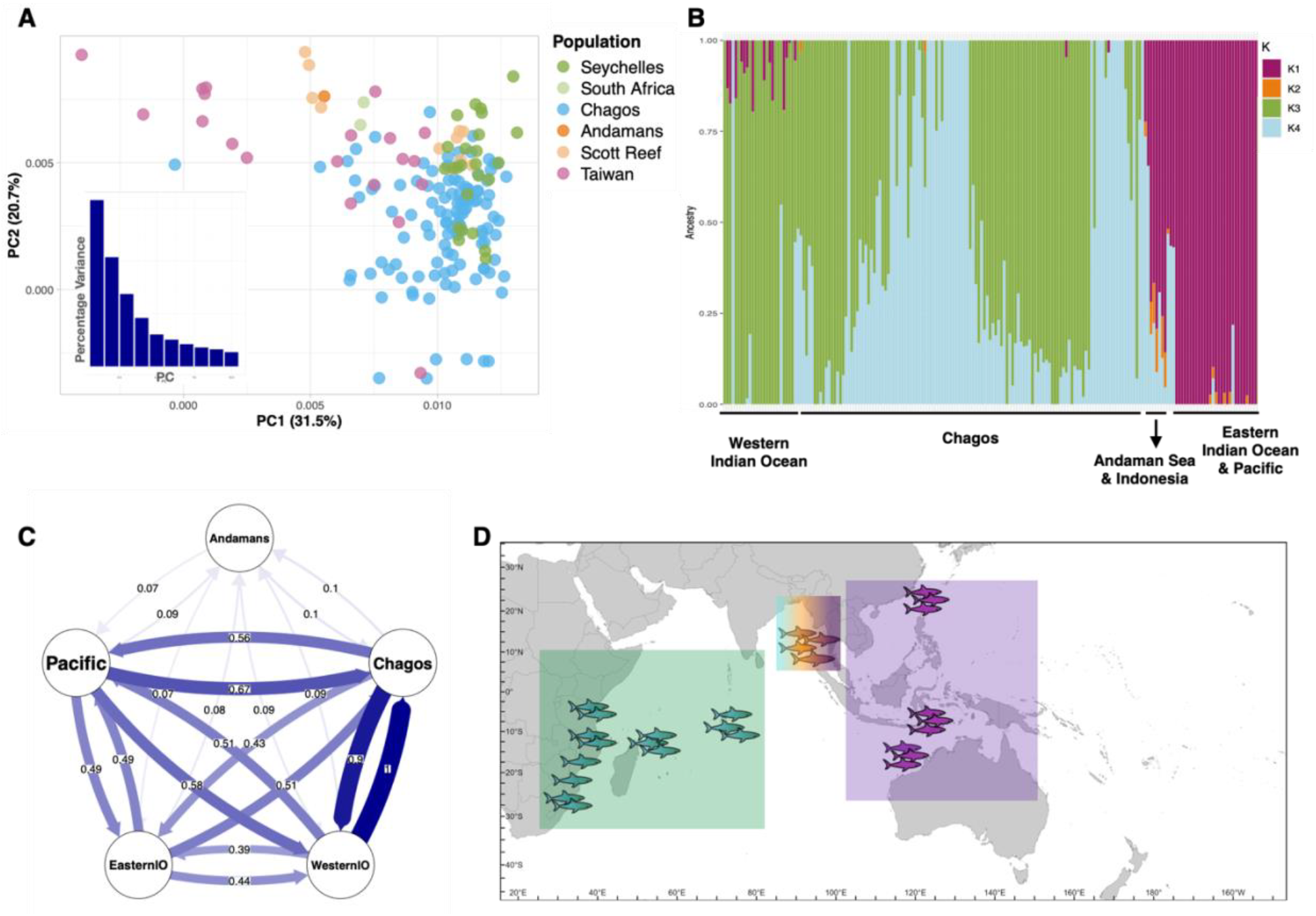
Population structure in silvertip sharks across the Indo-Pacific. (A) Principal component analysis (PCA) showing genetic differentiation among regional groups. (B) Admixture plot indicating ancestry proportions and evidence of genetic mixing between populations. (C) Directional migration and gene flow plot illustrating asymmetrical connectivity across regions. (D) Map of resulting sub-populations derived from these analyses, revealing two major genetic clusters (Western Indian Ocean and Chagos–Andaman–Eastern Indian Ocean–Pacific), which correspond to two distinct management units.

At K = 3, Chagos was resolved as a distinct cluster, and individuals from the Andamans, Indonesia, and Scott Reef showed signs of recent admixture between Chagos and EIO lineages (Figure 3B). At K = 2, populations grouped into two broad clusters: one comprising the EIO and Pacific, and the other including the WIO and Chagos (Supplementary Figure 3). At K = 4, additionally admixed individuals were detected, particularly between Chagos and the Andaman/Indonesia cluster (Supplementary Figure 3).

Genome-wide F_ST_ analyses revealed similar regional population structures to those observed in *C. amblyrhynchos*, though with lower overall differentiation (Supplementary Figure 4A, Supplementary Table 3 & 4). The Chagos population showed the greatest differentiation from the Pacific (F_ST_ = 0.009), followed by the EIO (F_ST_ = 0.006), and the Andaman region (F_ST_ = 0.007), while it was least differentiated from the WIO (F_ST_ = 0.005). The Andaman population exhibited the highest differentiation from other regions, whereas the EIO and Pacific populations were the least differentiated (F_ST_ = 0.001).

Although pairwise F_ST_ values were generally lower than in *C. amblyrhynchos*, regional trends were consistent in population wise comparisons (Supplementary Figure 4B). Populations within geographic clusters showed low differentiation, while F_ST_ values increased with geographic distances. WIO populations were most distinct from those in the Andaman and Indo-Pacific clusters.

As with grey reef sharks, we evaluated gene flow in *C. albimarginatus* using 10,000 SNPs from 195 individuals. Results showed significantly high gene flow between Chagos and the WIO, and lower levels of gene flow between all other regions, except the Andaman Sea, where no significant gene flow was detected (Figure 3C). Population group assignments, reflecting the major genetic clusters identified for the silvertip sharks, are shown in Figure 3D.

### Mitogenome Variation and Structure

Mtochondrial genome sequences, ~14,030 bp in length, were analyzed from 205 *Carcharhinus amblyrhynchos* individuals, along with one *Rhinoptera brasiliensis* used as an outgroup, to investigate phylogenetic relationships and haplotype sharing across the Indo-Pacific. A total of 1,040 variable sites (~7%) were identified, including 129 parsimony-informative positions, resulting in 19 distinct mtDNA haplotypes. Haplotypes, 1, 2 and 3 were the most dominant and originated in the Andaman Sea, eastern Indian Ocean (EIO). In the Pacific, haplotypes 4 and 5 predominated in the western Indian Ocean (EIO) and the Chagos archipelago (Supplementary Figure 5A). Nucleotide diversity across regional populations was π = 0.0031 (Supplementary Table 5), with a sample size of 223 and an estimated mutation rate of μ ≈3.74 × 10^−6^.

Phylogenetic reconstructions using both Median-joining networks and Bayesian inference revealed congruent topologies and well-supported clades, indicating spatially structured populations of *C. amblyrhynchos* (Figure 4A). Genetic distances between *C. amblyrhynchos* and the outgroup exceeded 25%. Among-region distances ranged from 0.3% to 8%, while within-region distances ranged from 0.1% to 4% in Chagos, 0.1% to 5% in the WIO, and 0.3% to 5% in the EIO. AMOVA results revealed significant genetic differentiation between regions (pairwise ΦST = 0.5647, *p* = 0.001), though pairwise ΦST values among individual populations were not significant. Tajima’s D values indicated no evidence of recent population expansion.

**Figure 4.**
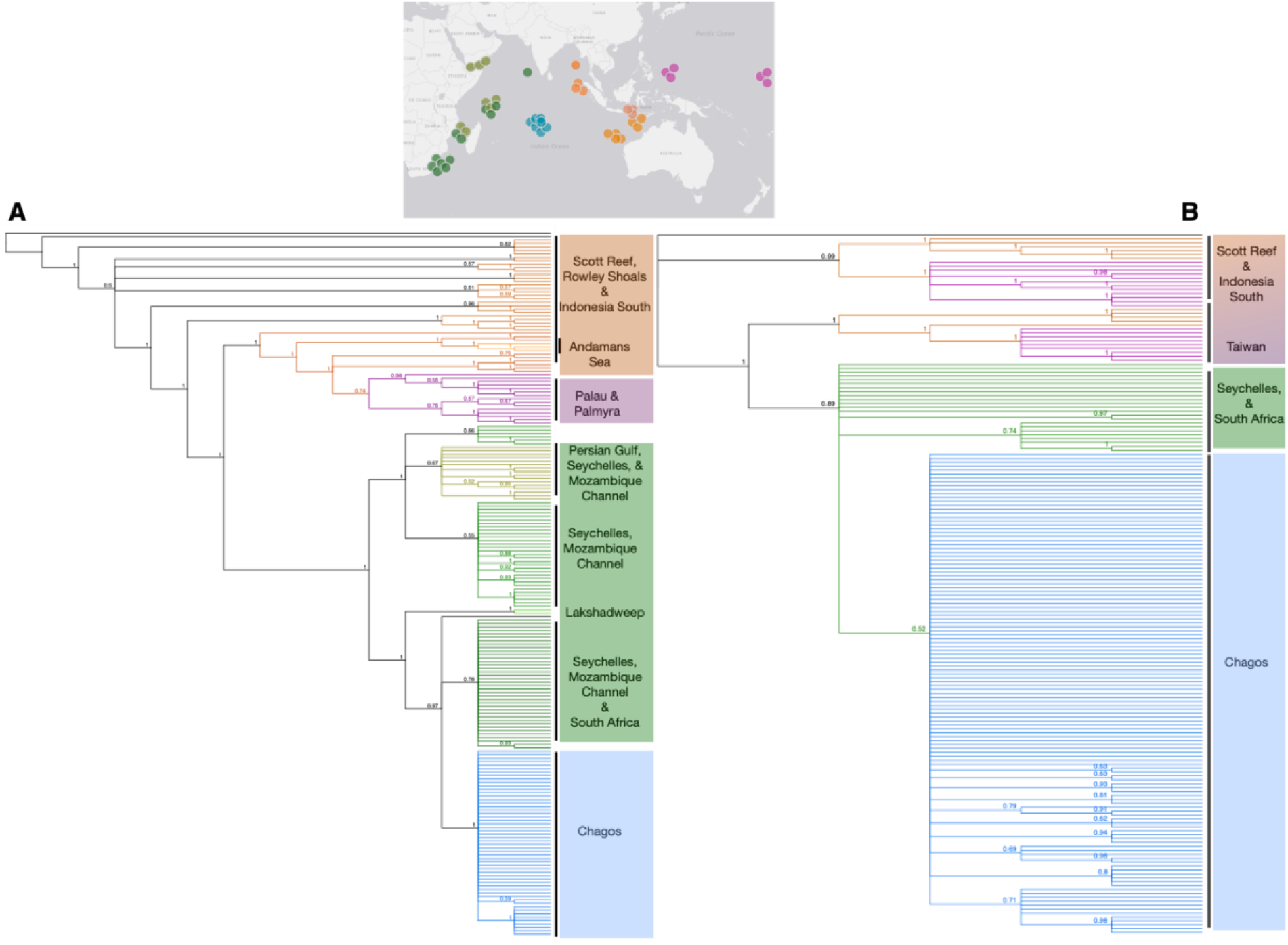
Mitochondrial genome phylogeny of reef sharks across the Indo-Pacific. (A) Grey reef sharks showing four distinct phylogenetic clusters: Western Indian Ocean (green), Chagos Archipelago (blue), Eastern Indian Ocean–Andaman Sea (orange), and Pacific (magenta). (B) Silvertip sharks showing three clusters: Western Indian Ocean (green), Chagos Archipelago (blue), and a combined Eastern Indian Ocean–Andaman Sea–Pacific group (orange/magenta). The map above indicates the geographic distribution of sampling sites, with colors corresponding to the clusters identified in the phylogenies.

For *Carcharhinus albimarginatus*, mitochondrial genomes (14,091 bp) were analyzed from 93 individuals and one *R. brasiliensis* outgroup. We identified 1,464 variable sites (~10%), including 102 parsimony-informative sites, and 13 haplotypes. Among the five main haplotypes 1 was predominant in the EIO, 2 in the Pacific, 3 in the EIO, Pacific and Chagos, 4 was frequent in the WIO and 5 was the main haplotype in the Chagos and upto a lesser extent in the WIO (Supplementary Figure 5B). Nucleotide diversity was lower than in *C. amblyrhynchos* (π = 0.0012, Supplementary Table 5), with a sample size of 195 and mutation rate μ≈1.54×10^−6^.

Phylogenetic analyses again revealed geographically structured clades with high bootstrap values and significant posterior probability support (Figure 4B). Genetic distances between *C. albimarginatus* and the outgroup exceeded 25%. Among-region distances ranged from 0.2% to 9%, and within-region distances ranged from 0.2% to 6% in Chagos, 0.1% to 5% in the WIO, and 1% to 7% in the EIO. AMOVA indicated low but significant ΦST values between regions and among populations. Neutrality tests (Tajima’s D) were non-significant, suggesting a lack of recent demographic expansion.

### Demographic History and Nucleotide Diversity

Analyses of demographic history in the grey reef shark populations using Pairwise Sequential Markovian Coalescence, suggested historical population sizes in the range of 200,000 to 110, 000 for the western Indian Ocean populations (Figure 6A, Supplementary Figure 6A), and 110,000 to 75, 000 in the Central Indian Ocean populations (Figure 6A, Supplementary Figure 6B) and the eastern Indian and Pacific Ocean populations (Figure 6A, Supplementary Figure 6C). All grey reef populations experienced a precipitous decline around 3-5 million years ago, with stabilization at a lower population level of ~30,000-50,000 individuals ~500, 000 years ago. All populations except for the eastern Indian Ocean and Pacific Ocean populations continued to decline beyond this point with a final stabilization at 10,000-15,000 individuals ~100,000 years ago (Figure 6A). The Eastern Indian Oceana and Pacific Ocean populations stabilized at around 30,000 individuals ~100,000 years ago (Figure 6A, Supplementary Figure 6C).

**Figure 6.**
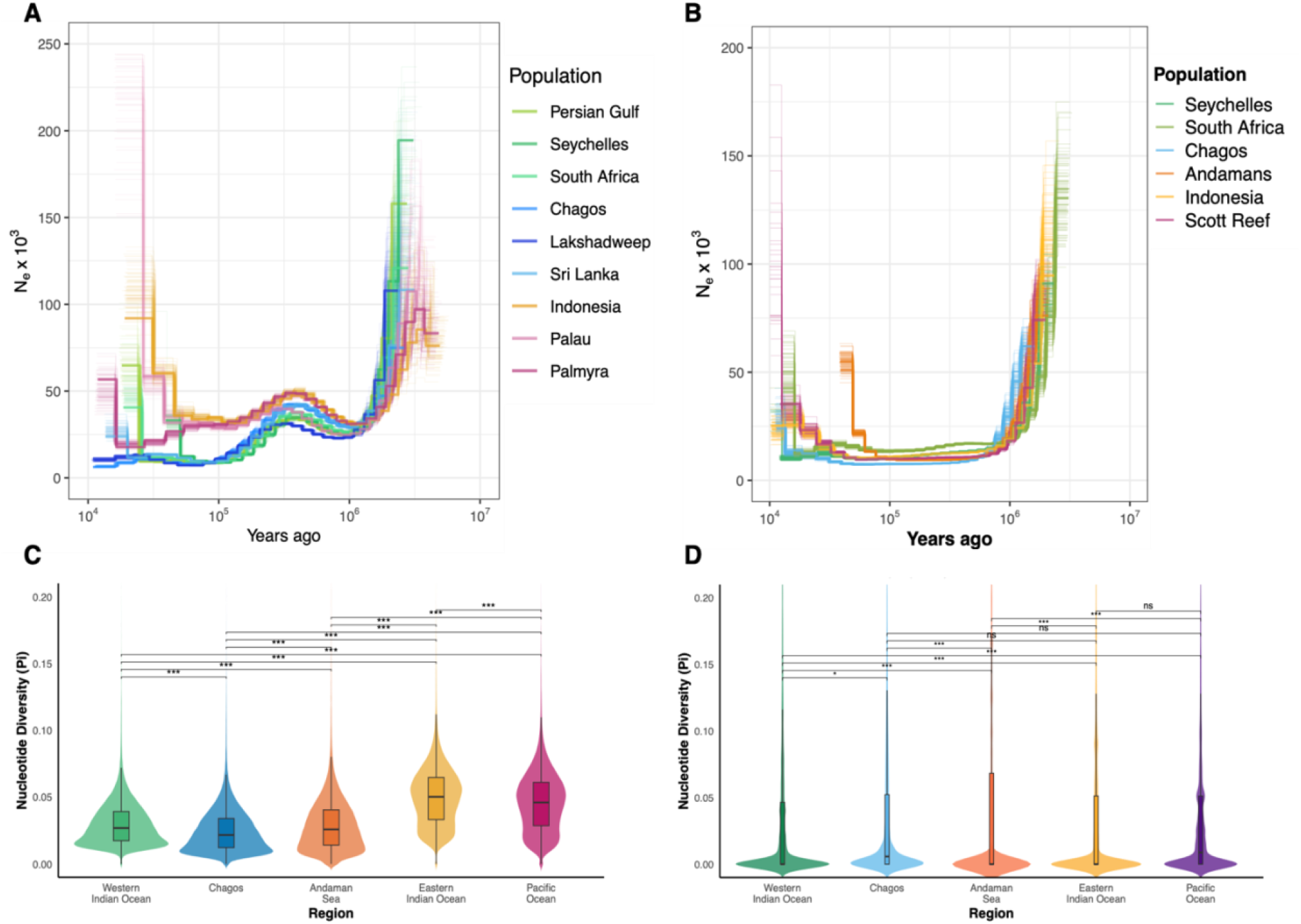
Demographic history and nucleotide diversity of reef shark populations. (A) Pairwise Sequentially Markovian Coalescent (PSMC) plot showing historical effective population size trajectories in grey reef sharks. (B) PSMC plot for silvertip sharks, illustrating contrasting demographic histories. (C) Nucleotide diversity across population groups of grey reef sharks. (D) Nucleotide diversity across population groups of silvertip sharks. Colors indicate population groups as shown in the figure legends.

Demographic history analyses in the silvertip shark populations using PSMC, revealed historical population sizes in the range of 130,000 to 90, 000 for the western Indian Ocean populations (Figure 6B, Supplementary Figure 6D), 60,000 for the Chagos population (Figure 6B, Supplementary Figure 6E) and 90,000 to 75, 000 in the Andaman Sea, and Eastern Indian Ocean populations (Figure 6B, Supplementary Figure 6F). Silvertip shark populations experienced a precipitous decline around 2-3.5 million years ago, with stabilization at ~15,000 individuals starting at ~750, 000 years ago (Figure 6B).

We estimated the average nucleotide diversity (π) within individuals of each population group for the grey reef and silvertip sharks. A total of 203833 neutral variant and non-variant SNP loci were used to calculate π for grey reef shark population groups and these resulted in π values in the range of 0.02 to 0.05 (Figure 6C, Supplementary Table 6). Each cluster was significantly different with respect to π and the EIO and Pacific Ocean clusters had the highest π values of ~0.05.

We used 289045 neutral variant and invariant SNP loci to calculate π for silvertip shark population groups. The π values were significantly lower compared to that for the grey reef shark populations, and most were in the range of 0.001 while the Chagos population had a π of 0.006 and the Pacific Ocean cluster had the highest π of 0.01 (Figure 6D, Supplementary Table 6). Distribution was skewed for all silvertip populations, with long whiskers and higher maximums suggesting a few individuals with elevated diversity. In a similar pattern to the grey reef sharks, the Pacific Ocean cluster of silvertips had a higher number of individuals with elevated diversity compared to other silvertip population groups.

### Tracking Shark Fisheries using Genomics

To evaluate the use of genomics to identify independent samples, a model was trained using supervised machine-learning to discriminate between population groups of the grey reef shark. Monte-Carlo cross-validation was utilized on the trained model to estimate the mean and variance of assignment of samples and accuracy was evaluated by resampling of random individuals from the training dataset. Overall assignment accuracy of the model for the training data was greater than 93% for all population groups when 95% of individuals in each population and all loci from the training data set were utilized(Figure 7A). Assignment accuracy was greater than 93% for the Western IO, greater than 96% for the Chagos, and ~100% for the Easter IO and Pacific Ocean populations (Figure 8A).

**Figure 7.**
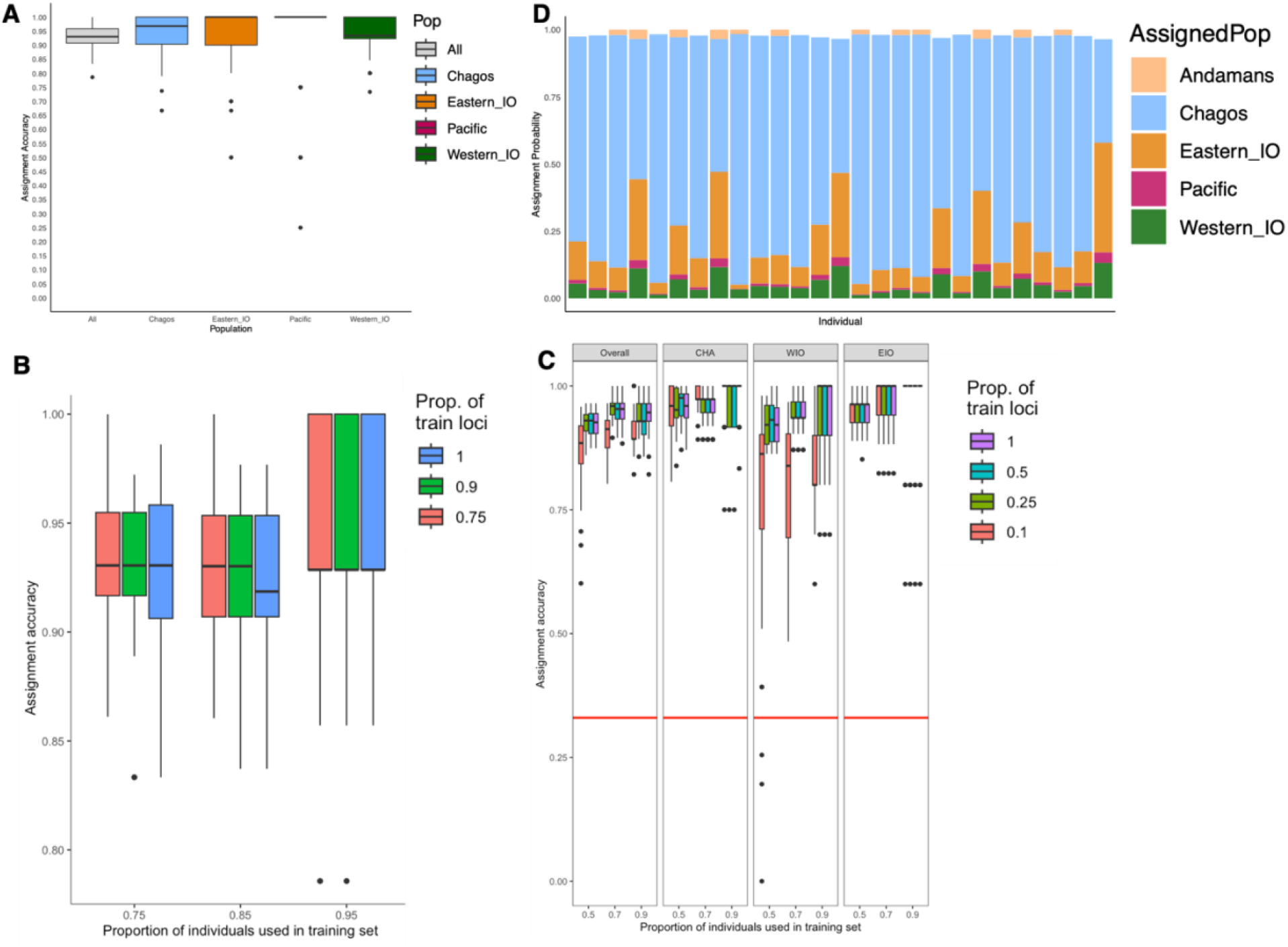
Tracking fisheries using the Reefshark Genomescape. (A) Overall assignment accuracy of grey reef shark reference populations. (B) Assignment accuracy across different proportions of individuals and numbers of training loci. (C) Assignment accuracy per population group under varying proportions of individuals and training loci. (D) Assignment probabilities of fished sharks sampled in the Chagos Archipelago.

The overall assignment accuracy was tested with varying proportions of individuals and loci from the training dataset, to assess the accuracy of the model when using smaller datasets. The model achieved median assignment accuracy of greater than 93% when the proportion of individuals was changed between 75%, 85% or 95% of individuals in each population group and the proportion of training loci was changed between 75%, 90% and 100% of the total loci for each population (Figure 8B). However, when 95% of individuals from each population were used in the training dataset the variance of accuracy skewed towards 100% accuracy, suggesting a higher probability of accurate results when 95% or higher percentage of individuals in the training dataset are used (Figure 8B).

We then tested the impact of using smaller proportions of training data with respect to identifying individuals and loci for assignment accuracy of individual population groups. Proportions of individuals used were 50%, 70% or 90% and proportion of loci varied between 10%, 25%, 50% and 100%. Overall median assignment accuracy was at 85% or higher even when 50% of individuals and 10% of training loci were used (Figure 8C). Cumulative assignment accuracy as well as assignment accuracy for each individual population group was most optimal when 90% of the individuals in each population were used, and 25% or higher proportion of training loci were used (Figure 8C). In summary, the number of individuals had a greater impact on assignment accuracy than the proportion of loci, when loci above a threshold of 25% were used.

To evaluate the discriminatory power of this model in assigning unknown samples (those not included in the training data and never previously encountered by the model) we tested this using genetic data from 27 samples confiscated from a vessel illegally fishing in the Chagos archipelago. These samples were treated as unassigned during the assignment test. The model correctly assigned 26 out of 27 samples (96%) to Chagos, based on assignment probabilities greater than 50% for the region (Figure 8D). One of the 27 samples had a less than 50% assignment probability for the Chagos and therefore it was not assigned to the Chagos grey reef shark population.

Admixture analyses of the confiscated grey reef shark samples from the Chagos Archipelago grouped them with the local Chagos population, supporting our assignment based on a parallel analysis discussed above (Supplementary Figure 5A).

In contrast, the silvertip shark samples were assigned to the Western Indian Ocean population (Supplementary Figure 5B). However, high gene flow, due to likely due to migration between the Western Indian Ocean and Chagos, makes it difficult to confidently assign these individuals to a specific population.

## DISCUSSION

This study presents the most comprehensive population genomic analysis to date of two ecologically significant reef sharks—the grey reef shark (*Carcharhinus amblyrhynchos*) and the silvertip shark (*C. albimarginatus*), across their Indo-Pacific range. Our findings reveal contrasting patterns of genetic diversity, population structure, and demography, including novel patterns of connectivity and isolation that point to underlying oceanographic processes warranting further investigation. Crucially, this work also establishes a robust genomic framework for fisheries traceability and conservation enforcement in one of the world’s most heavily fished oceans, providing an urgently needed tool for safeguarding reef sharks and the ecosystems they structure.

### Patterns of Genetic Diversity, Connectivity and Demography

Our results reveal strong genetic structuring in grey reef sharks, with populations divided into four major clusters: Western Indian Ocean, Chagos Archipelago, Andaman Sea, and a combined Eastern Indian Ocean -Pacific group. The Andaman populations were nearly completely isolated, showing negligible gene flow with neighboring regions. Silvertip sharks, by contrast, show weaker structuring and higher connectivity, particularly between the WIO and the Chagos and EIO and the Pacific, though the Andaman populations remained distinct.

These contrasting patterns are consistent with telemetry observations: grey reef sharks are site-attached with limited movement ranges, while silvertips are wider ranging and more dynamic (Jacoby et al. 2020; Williamson et al. 2021; Daly et al. 2023; Dwyer et al. 2020). Connectivity patterns broadly follow biogeographic provinces but also reveal notable contrasts, including unexpectedly high connectivity between the Chagos Archipelago and the Western Indian Ocean, and near-complete isolation of populations in the Andaman Sea. Together, these findings demonstrate how historical demography, and ecological barriers, have shaped the genetic landscape of both species.

The connectivity patterns we observed, appear to be shaped by oceanographic processes. For instance, the Chagos–WIO connection between grey reef sharks is surprising because previous studies tracking their movements showed relatively limited dispersal within the region (Daly et al. 2023). However, our observation parallels green turtle migration routes driven by the South Equatorial Current (Hays et al. 2014), while whale sharks similarly track productivity fronts across the Indian Ocean (Arrowsmith et al. 2021). In contrast, the near complete isolation of Andaman Sea populations suggests an oceanographic break limiting dispersal, a barrier also reported in marine turtles (Wallace et al. 2010). These observation across multiple species warrant further studies, including genetic as well as tracking studies of a higher number of individuals per species to clarify mechanisms constraining gene flow.

The Indian Ocean remains one of the least-studied regions regarding shark connectivity and migration. Complex transboundary movements within the WIO (Barkley et al. 2019), along with extended residency and movement corridors in other parts of the Indian Ocean (Sequeira et al. 2025), suggest that large-scale connectivity patterns are still poorly understood. Connectivity between shark populations in the WIO and Chagos, in particular, has not been fully quantified. Further research integrating genetic sampling, telemetry, and ecological data—including age, migration timing, oceanographic conditions, prey availability, and key behaviors such as foraging and mating—is needed to clarify the mechanisms constraining gene flow and shaping population structure across this region.

From a conservation perspective, these results underscore the need for species-specific strategies. Grey reef sharks comprise multiple genetically isolated subpopulations that function as discrete management units. Given their restricted movement ranges and high site fidelity, effective conservation will require localized protection, ideally through the establishment and enforcement of marine protected areas (MPAs) as suggested by other studies of movement patterns as well (Dwyer et al. 2020). Although these units span multiple national jurisdictions, their limited dispersal capacity and low transboundary movements (Daly et al. 2023) suggests that well-managed local or cross-boundary MPAs could provide meaningful protection if effectively enforced. Silvertip sharks, in contrast, require coordinated cross-jurisdictional measures that extend across open-ocean habitats. Their broader dispersal, long-distance movements, and evidence of transboundary—and in some cases transoceanic—population connectivity necessitate management frameworks that operate beyond national waters, through strengthened regional fisheries agreements and cooperative international governance.

Our analyses also reveal contrasting demographic trajectories in the two species, after they experienced similar demographic contractions 2–5 million years ago, during the Pleistocene, similar to many coastal species (Ludt and Rocha 2015; Tillett et al. 2012). Grey reef sharks exhibit relatively elevated nucleotide diversity and effective population sizes, particularly in the EIO and Pacific, consistent with more stable demographic histories and potential persistence within climatic refugia in the Pacific during Pleistocene glacial cycles. The stabilization of grey reef sharks at higher effective sizes in the Pacific suggests a demographic pattern comparable to that of sperm whales, which are thought to have persisted in and diversified from the Pacific as a climatic refugium (Morin et al. 2018). These patterns suggest that Pacific reef systems may have buffered grey reef sharks against severe declines, while populations in the WIO and Chagos underwent sharper reductions. Silvertip sharks, by contrast, exhibit markedly lower genetic diversity across most of their range, indicative of reduced effective population sizes and inbreeding. In addition to historical declines associated with geological events, the species’ large body size and frequent capture in coral reef fisheries suggest that recent population reductions are also likely driven by contemporary fishing pressures. The consistently low genetic diversity in silvertip sharks indicates reduced adaptive potential to environmental change, small effective population sizes, and heightened vulnerability to fishing pressure. These findings indicate that the current IUCN listing of silvertip sharks as Vulnerable (Rigby, C.L. et al. 2023) likely underestimates their extinction risk. Our results support an uplisting to Endangered, given their severe population declines and low genetic diversity, which may render some populations functionally extinct.

### Conservation and Policy Implications

The loss of large-bodied predators such as reef sharks disrupts trophic pyramids, triggering cascades including mesopredator release, loss of key herbivores, and widespread reef degradation(Graham et al. 2017; Roff et al. 2016; Bornatowski et al. 2014). These changes reduce ecosystem resilience and compromise the services reefs provide. Coastal and Indigenous communities that rely on coral reefs for food security and livelihoods are particularly affected, as ecosystem imbalances threaten fisheries and exacerbate vulnerability to overexploitation and environmental change (Cisneros-Montemayor et al. 2016; Eddy et al. 2021).

Despite global efforts to curb illegal, unreported, and unregulated fishing, enforcement is hampered by technical, legal, and geopolitical loopholes, including transshipment, and flag-hopping, allowing “dark vessels,” to operate without detection (Belhabib and Le Billon 2020;Belhabib and Le Billon 2022; Paolo et al. 2024). These gaps undermine sustainable fisheries governance, threaten biodiversity, and harm the livelihoods of lawful fishers (Okafor-Yarwood et al. 2022).

Beyond advancing ecological understanding, this study establishes a practical enforcement framework. The Reefshark Genomescape, the largest genomic reference database and spatially explicit map of intraspecies genetic diversity for any reef shark, provides a scalable genomic tool to overcome key limitations in fisheries monitoring and conservation enforcement. As aspatially explicit map of intraspecific diversity for two reef shark species, it enables assignment of fished individuals to their source populations across the Indo-Pacific. The strong genetic structure in grey reef sharks allows high-resolution assignment to specific sub-populations, and in some cases, to individual local populations. In contrast, the higher connectivity of silvertip sharks supports assignment to broader sub-populations that span ocean basins. Both species are frequently targeted by fisheries (Sherman et al. 2023) and can serve as biological indicators of fishing pressure. Grey reef sharks, in particular, by virtue of their strong population structure, extremely small movement corridors and low transboundary crossings (Daly et al. 2023) and frequent capture in fisheries (Bennett et al. 2022), can be used to determine the geographic origins of entire fisheries catch consignments, as well as in the broader trade in derivative products (e.g., shark fins, meat). Integrated into fisheries and trade monitoring, this framework will enable identification of fishing hotspots, detection of IUU activity, and guidance for spatially explicit Fisheries Management Zones (FMZs). Comparable genomic approaches in terrestrial conservation, such as tracking poaching and trade of African elephants, have demonstrated the transformative potential of this methodology (Wasser et al. 2015; Wasser et al. 2018).

The Indo-Pacific, particularly the Indian Ocean, is both a biodiversity hotspot, and a globally exploited region for sharks and rays (Dulvy et al. 2014; Dulvy et al. 2021; Dent and Clarke 2015; Jabado et al. 2018; Pollom et al. 2024). Both species face some of their greatest threats within this region, underscoring the need for conservation measures across their full Indo-Pacific distribution to safeguard species populations (Bennett et al. 2022; Jabado et al. 2017).Genomic traceability provides incorruptible evidence linking shark products to geographic origin, bypassing enforcement gaps, strengthening CITES Appendix II compliance, and supporting regional fisheries agreements. By integrating ecological insights with practical conservation tools and regular genetic monitoring, the Reefshark Genomescape provides a scalable, adaptive framework to track fishing pressure, guide fisheries regulation, enable targeted enforcement, and safeguard reef shark populations, and, by extension, the reef ecosystems they structure, in heavily exploited oceanic regions.

## CONCLUSION

This study establishes the Reefshark Genomescape as both a foundational scientific resource for understanding patterns of genetic diversity and the ecological processes driving them, and as a practical tool for ocean surveillance and conservation enforcement. It represents the first range-wide genomic framework for ecological monitoring, connectivity assessment, and management of two ecologically critical reef sharks across the Indo-Pacific, and, by extension, the reef ecosystems they support.

We identified both isolated and connected subpopulations, highlighting vulnerabilities to overfishing and environmental change. By enabling forensic tracing of shark products to their source populations, the framework provides a scalable approach to identify ecosystems under pressure, guide targeted management, inform spatially explicit fisheries regulations, and enforce international trade restrictions. In an era of accelerating biodiversity loss, this study serves as a model for other marine species and underscores the essential role of genetic metrics in long-term, adaptive conservation strategies.

Looking forward, this work lays the foundation for investigating the ecological processes structuring populations in the Indo-Pacific. Integrating range-wide genomic data with oceanographic modeling and telemetry will clarify the mechanisms underlying connectivity and isolation. Applying these insights to current management frameworks will support the design of effective marine protected areas, cross-boundary regulations, and ecosystem-scale conservation strategies. The Reefshark Genomescape provides a scalable, adaptive model for safeguarding apex predators and the reef ecosystems they structure in one of the world’s most heavily exploited oceans.

## Supporting information

Link text is included in the supplemental figures and tables..

## FUNDING SOURCES

We thank the Bertarelli Foundation for providing the grant which supported this project. We thank Biology and Oceans Departments at Stanford University for institutional, computational and administrative support of the project.

## AUTHOR CONTRIBUTIONS

Conceptualization, and study design: SJ with input from all authors., Project Advisory role: BB, Sampling: SJ co-ordinated the sampling by co-authors across study area and conducted sampling in Lakshadweep and Seychelles. GA coordinated sampling and shipping of samples from Seychelles, Mozambique Channel and Indonesia. RS, BB collected samples in the Chagos archipelago. All authors contributed to fieldwork in respective locations, data collection, and data formatting. Methodology and Analyses: SJ. Visualization: SJ. Funding acquisition: SJ. and BB. Project administration: SJ. Writing – original draft: SJ. Writing – review & editing: SJ., BB., GA., RJ., JN., and RD.

## ACKNOWLEDGEMENTS

Thanks to the Administração Nacional das Áreas de Conser-vação (ANAC), Mozambique, and Instituto Oceanográfico de Moçambique (InOM) for permitting and supporting ongoing research in Mozambique. Thanks to the iSimangaliso Marine Protected Area and iSimangaliso Wetland Park management and Ezemvelo KZN Wildlife, together with the Maputo National Park and management, for their ongoing support of this research.

## CONFLICTS OF INTEREST

The authors declare no conflicts of interest.

## Notes

### Competing Interest Statement

The authors have declared no competing interest.

### Summary of Updates

The manuscript has been revised to reflect editing of the written text and reorganization of the figures to improve clarity and readability with inputs from co-authors of the manuscript.

