## Supplementary material for "Mapping Risk and Conservation Potential Across the Indo-Pacific with Reefshark Genomescapes": Link text is included in the supplemental figures and tables..

### Supplementary Figures and Tables

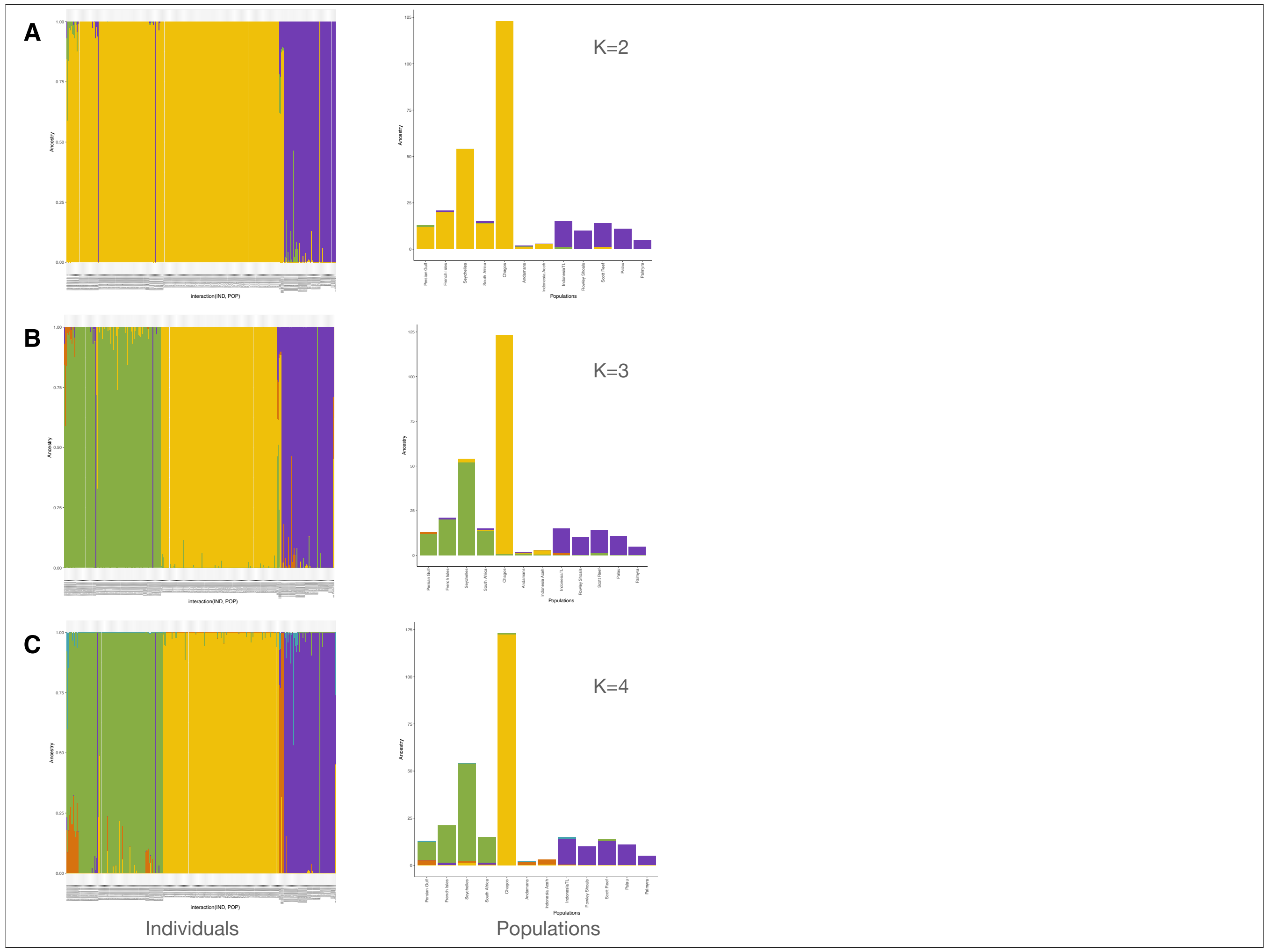


**Supplementary Figure 1.** ADMIXTURE analysis of grey reef sharks at K = 2, 3 and 4.
Bar plots show individual ancestry proportions, illustrating how population structure becomes increasingly resolved with higher K values.

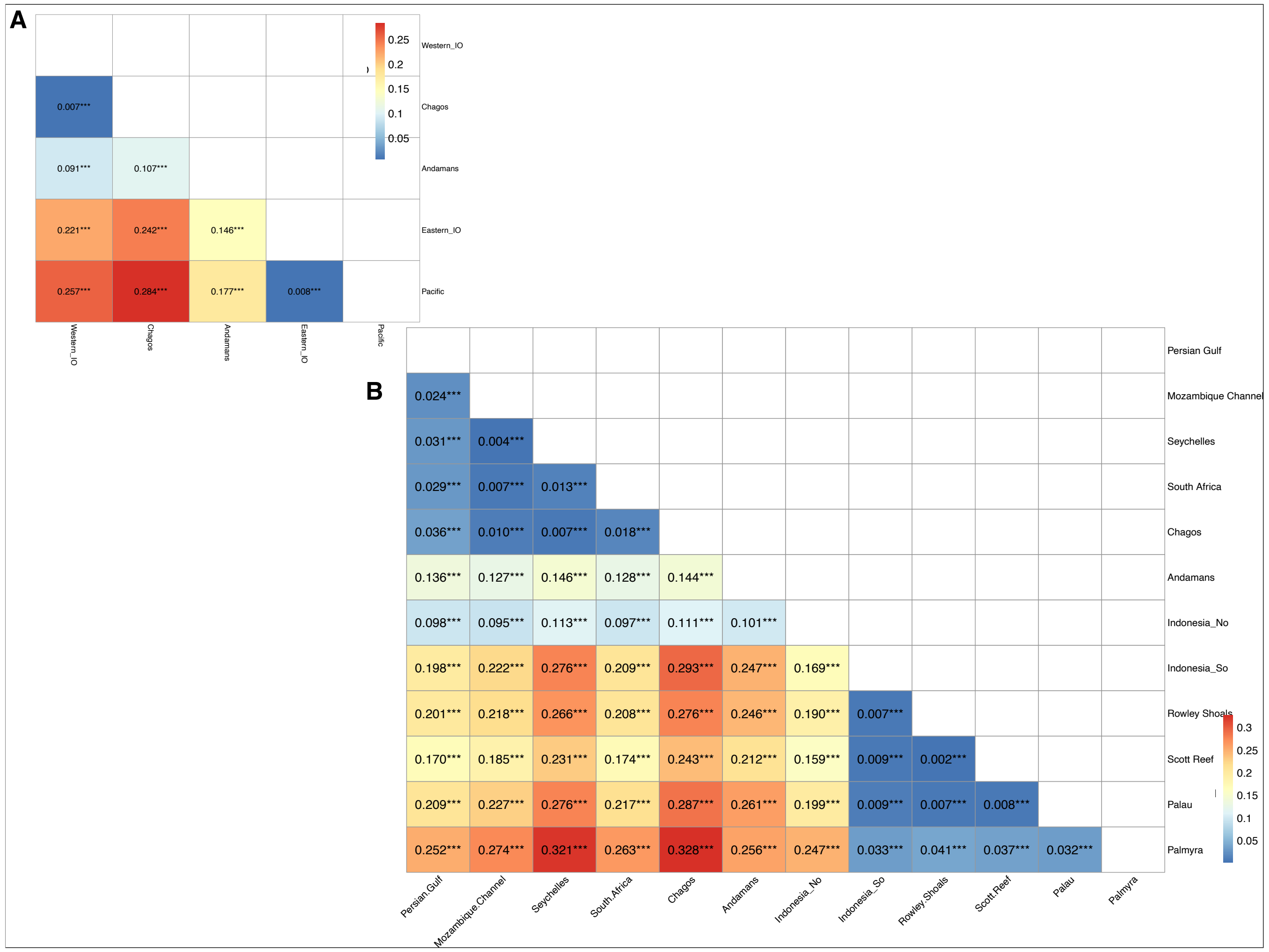


**Supplementary Figure 2**: Fst matrix by population group and by individual population for grey reef sharks

#### Supplementary Table 1: *C. amblyrhynchos* group pairwise FST_matrix

| GRSgroup_pairwise_FST_matrix | | | | | |
|---|---|---|---|---|---|
| | **Western_IO** | **Chagos** | **Andaman Sea** | **Eastern_IO** | **Pacific** |
| **Western_IO** | NA | NA | NA | NA | NA |
| **Chagos** | 0.00728 | NA | NA | NA | NA |
| **Andamans** | 0.09088 | 0.10657 | NA | NA | NA |
| **Eastern_IO** | 0.22129 | 0.24241 | 0.14583 | NA | NA |
| **Pacific** | 0.25748 | 0.28353 | 0.17735 | 0.00819 | NA |

#### Supplementary Table 2: *C. amblyrhynchos* population pairwise FST_matrix

| GRSpopulation_pairwise_FST_matrix | | | | | | | | | | | | |
|---|---|---|---|---|---|---|---|---|---|---|---|---|
| | **Chagos** | **South Africa** | **Scott Reef** | **Palmyra** | **Palau** | **Persian Gulf** | **Rowley Shoals** | **Mozambique Channel** | **Seychelles** | **Andamans** | **Indonesia_So** | **Indonesia_No** |
| **Chagos** | NA | NA | NA | NA | NA | NA | NA | NA | NA | NA | NA | NA |
| **South Africa** | 0.0181 | NA | NA | NA | NA | NA | NA | NA | NA | NA | NA | NA |
| **Scott Reef** | 0.2428 | 0.17412 | NA | NA | NA | NA | NA | NA | NA | NA | NA | NA |
| **Palmyra** | 0.3277 | 0.26291 | 0.03692 | NA | NA | NA | NA | NA | NA | NA | NA | NA |
| **Palau** | 0.2872 | 0.21657 | 0.00788 | 0.0322 | NA | NA | NA | NA | NA | NA | NA | NA |
| **Persian Gulf** | 0.0358 | 0.02873 | 0.17049 | 0.25163 | 0.2095 | NA | NA | NA | NA | NA | NA | NA |
| **Rowley Shoals** | 0.2764 | 0.20766 | 0.00224 | 0.04079 | 0.0074 | 0.20089 | NA | NA | NA | NA | NA | NA |
| **Mozambique Channel** | 0.0098 | 0.0075 | 0.18482 | 0.27362 | 0.2273 | 0.02426 | 0.21796 | NA | NA | NA | NA | NA |
| **Seychelles** | 0.0072 | 0.01293 | 0.23094 | 0.32136 | 0.2763 | 0.03112 | 0.26639 | 0.00406 | NA | NA | NA | NA |
| **Andamans** | 0.144 | 0.12763 | 0.21212 | 0.25638 | 0.2611 | 0.13577 | 0.24558 | 0.12728 | 0.14611 | NA | NA | NA |
| **Indonesia_So** | 0.2932 | 0.20885 | 0.0087 | 0.03284 | 0.0093 | 0.19834 | 0.00654 | 0.22244 | 0.27608 | 0.24675 | NA | NA |
| **Indonesia_No** | 0.1111 | 0.09744 | 0.15937 | 0.24716 | 0.1987 | 0.09838 | 0.18963 | 0.09535 | 0.11309 | 0.1008 | 0.16856 | NA |

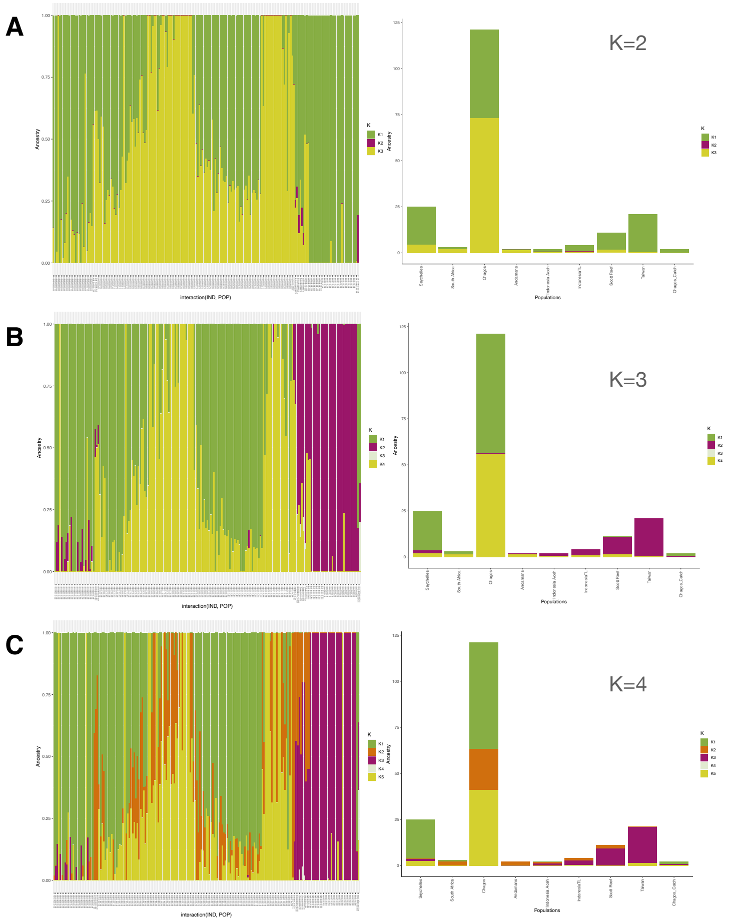


**Supplementary Figure 5: Admixture_ Silvertip Sharks**

**Supplementary Figure 3.** ADMIXTURE analysis of silvertip sharks at K = 2, 3 and 4.

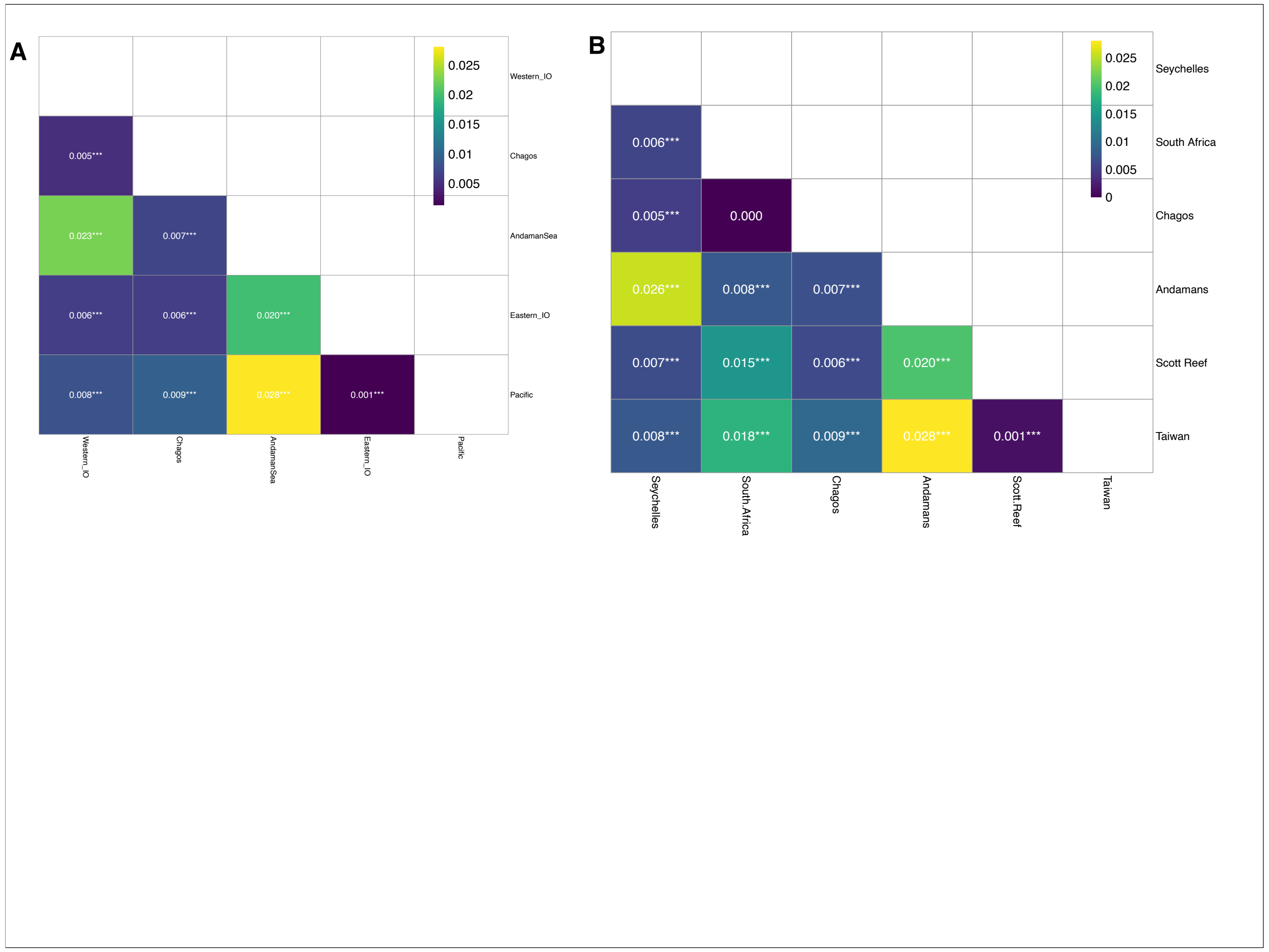


**Supplementary Figure 4**: Fst matrix by population group and by individual population for silvertip sharks

**Supplementary Table 3: *C. albimarginatus* group pairwise FST_matrix**

| STSgroup_pairwise_FST_matrix | | | | | |
|---|---|---|---|---|---|
| | **Chagos** | **Western _IO** | **Andama n [illegible]** | **Eastern_ IO** | **Pacific** |
| **Chagos** | NA | NA | NA | NA | NA |
| **Western _IO** | 0.00495 | NA | NA | NA | NA |
| **Andama n [illegible]** | 0.00685 | 0.02264 | NA | NA | NA |
| **Eastern_ IO** | 0.00632 | 0.00635 | 0.02003 | NA | NA |
| **Pacific** | 0.0094 | 0.00805 | 0.02808 | 0.00132 | NA |

**Supplementary Table 4: *C. albimarginatus* population pairwise FST_matrix**

| STSpopulation_pairwise_FST_matrix | | | | | | |
|---|---|---|---|---|---|---|
| | **Seychelles** | **South Africa** | **Chagos** | **Andamans** | **Scott Reef** | **Taiwan** |
| **Seychelles** | NA | NA | NA | NA | NA | NA |
| **South Africa** | 0.01637 | NA | NA | NA | NA | NA |
| **Chagos** | 0.00521 | 0.00498 | NA | NA | NA | NA |
| **Andamans** | 0.02582 | 0.02289 | 0.0069 | NA | NA | NA |
| **Scott Reef** | 0.00673 | 0.02189 | 0.0063 | 0.02003 | NA | NA |
| **Taiwan** | 0.00779 | 0.02696 | 0.0094 | 0.02808 | 0.00132 | NA |

**Supplementary Table 5: Mitochondrial Haplotype Network Statistics**

| Species | Variable Sites | Parsimony informative sites | Number of haplotypes | Nucleotide diversity (π) | Phi_ST | Pvalue | Tajima's D |
|---|---|---|---|---|---|---|---|
| *C. amblyrhynchos* | 1040 | 129 | 19 | 0.00306594 | 0.56472 | 0.001 *** | NS |
| *C. albimarginatus* | 1464 | 102 | 13 | 0.0012143 | NA | <0.001 *** | NS |

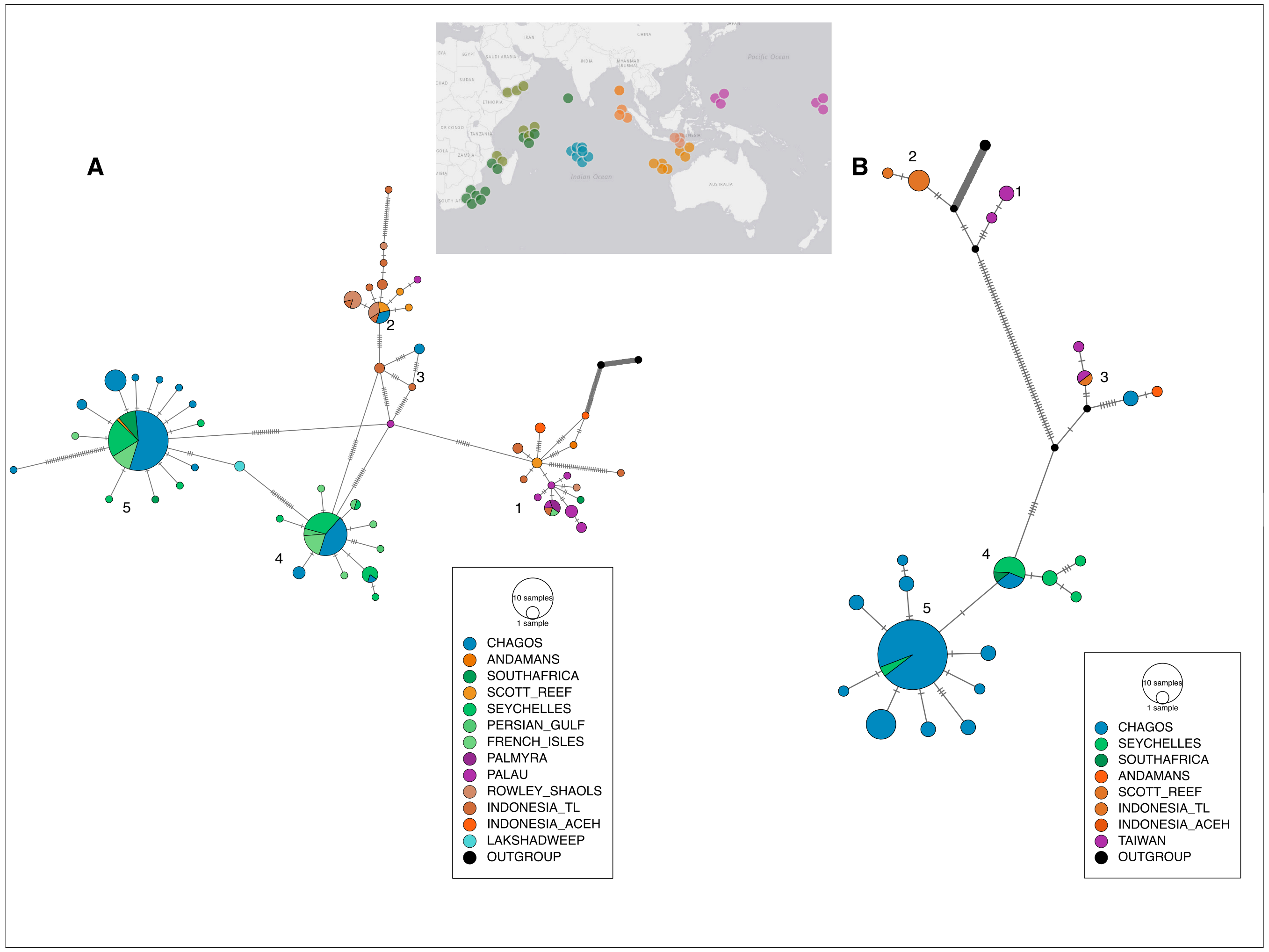


**Supplementary Figure 5. Mitochondrial haplotype networks of reef sharks.**
(A) Grey reef sharks, showing four major haplogroups corresponding to the Western Indian Ocean, Chagos Archipelago, Eastern Indian Ocean–Andaman, and Pacific.
(B) Silvertip sharks, showing five haplogroups corresponding to the Chagos Archipelago, Western Indian Ocean, Eastern Indian Ocean–Andaman–Pacific combined, additional unique haplotypes in the the Pacific and Eastern IO.

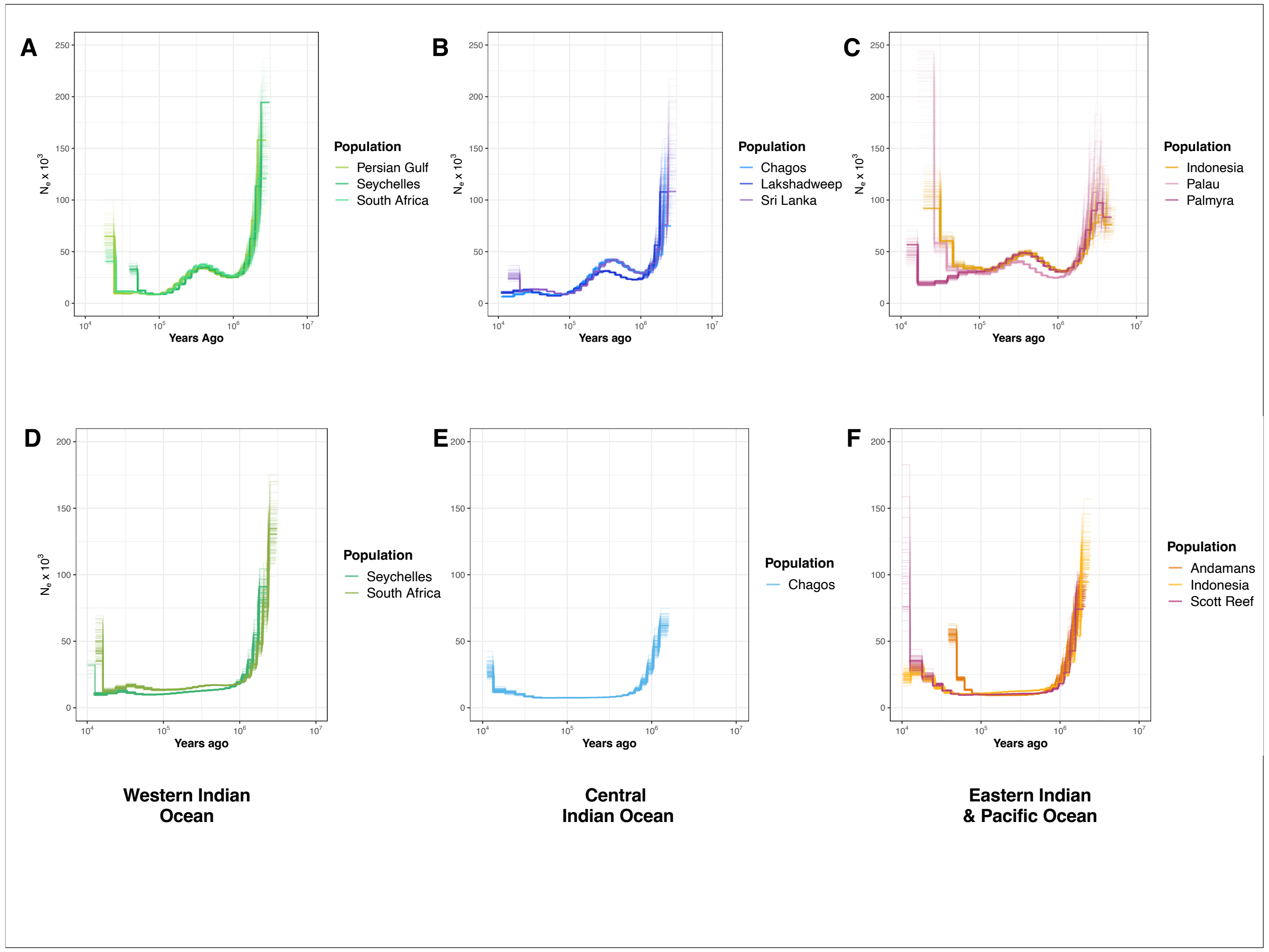


**Supplementary Figure 6. Pairwise Sequentially Markovian Coalescent (PSMC) plots of reef shark populations.**
(A–C) Grey reef shark populations from the Western Indian Ocean, Central Indian Ocean (Chagos/Andaman), and Pacific, respectively.
(D–F) Silvertip shark populations from the Western Indian Ocean, Central Indian Ocean (Chagos/Andaman), and Pacific, respectively.
Colors correspond to population groups as indicated in the legend.

#### Supplementary Table 6: Nucleotide Diversity

| GRSavg_pi_per_grp | | | | STSavg_pi_per_grp | | |
|---|---|---|---|---|---|---|
| **pop** | **mean_pi** | **number of sites** | | **pop** | **mean_pi** | **number of sites** |
| **Western Indian Ocean** | 0.03054663 | 203833 | | **Western Indian Ocean** | 0.001 | 289045 |
| **Chagos** | 0.02541538 | 203833 | | **Chagos** | 0.0064 | 289045 |
| **Andaman Sea** | 0.03000888 | 203833 | | **Andaman Sea** | 0.001 | 289045 |
| **Eastern Indian Ocean** | 0.05084639 | 203833 | | **Eastern Indian Ocean** | 0.001 | 289045 |
| **Pacific Ocean** | 0.04674426 | 203833 | | **Pacific Ocean** | 0.01 | 289045 |

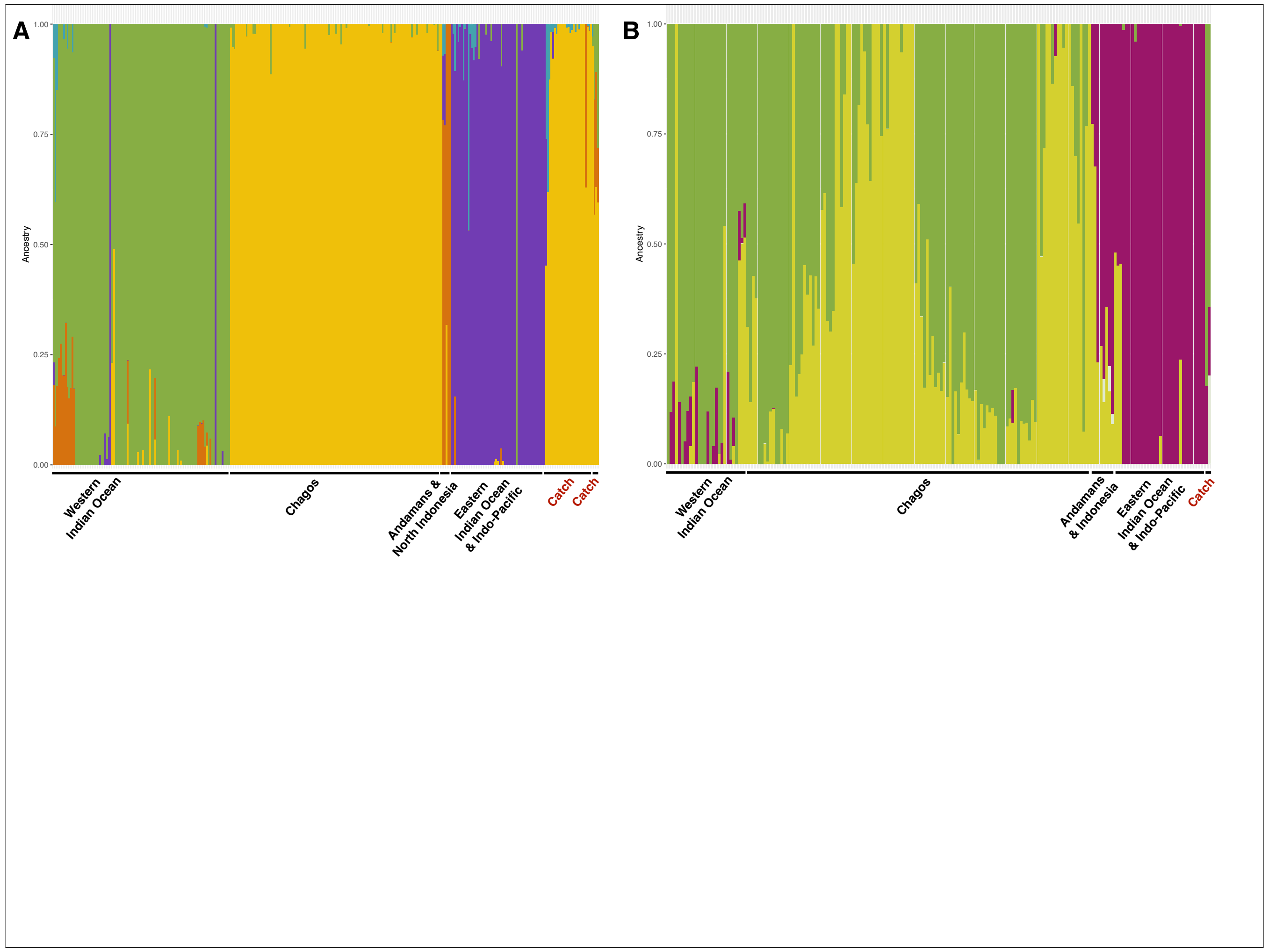


**Supplementary Figure 7. Tracking fisheries using admixture analyses.**
(A) Admixture analyses of grey reef sharks showing assignment probabilities of individuals to reference populations.
(B) Admixture analyses of silvertip sharks showing assignment probabilities of fished individuals to reference populations.
